# TAF1-dependent transcriptional dysregulation underlies multiple sclerosis

**DOI:** 10.1101/2024.08.23.609325

**Authors:** Claudia Rodríguez-López, Ivó H. Hernández, José Terrón-Bautista, Eneritz Agirre, Julia Pose-Utrilla, David Lozano-Muñoz, Irene Ruiz-Blas, Alejandra M. Arroyo, Inés García-Ortiz, Miriam Lucas-Santamaría, María González-Bermejo, María C. Ortega, Isabel Machín-Díaz, Juan C. Chara, Tjado H.J. Morrema, Chao Zheng, Zara Martínez, Fernando Pérez-Cerdá, Miriam Martínez-Jiménez, Beatriz Sancho-González, Alberto Pérez-Samartín, Mukund Kabbe, Marcos Casado-Barbero, María Santos-Galindo, Aldo Borroto, Balbino Alarcón, Alberto Paradela, Fernando de Castro, Nina L. Fransen, Jeroen J.M. Hoozemans, Claudio Toma, Carlos Matute, Diego Clemente, Felipe Cortés-Ledesma, Gonçalo Castelo-Branco, José J. Lucas

## Abstract

A major conceptual and clinical challenge in multiple sclerosis (MS) is understanding the mechanisms that drive the central nervous system (CNS)-resident neuroinflammation and neurodegeneration underneath disease progression. Genome-wide association studies (GWAS) have implicated RNA polymerase II (RNAPII) promoter-proximal pausing in oligodendrocyte pathology, but the causal mechanisms remain unclear. Here we find that the C-terminal region of TAF1, a core component of the general transcription factor TFIID, is underdetected in progressive MS brains, which can be explained by endoproteolysis due to extralysosomal cathepsin B (CTSB). Mice lacking the C-terminal TAF1 domain (Taf1d38) exhibit MS-like brain transcriptomic signature, alongside CNS-resident inflammation, progressive demyelination, and motor disability. Mechanistically, C-terminal TAF1 interacts with MS-linked factors that cooperate to regulate RNAPII pausing, particularly affecting oligodendroglial myelination genes. These findings uncover a previously unrecognized transcriptional mechanism underlying MS progression and establish a tractable *in vivo* model for therapeutic development.

## Introduction

MS is a disabling disease due to neuroinflammation and demyelination that affects 2.8 million people worldwide ^1,2^. Most individuals debut with relapsing–remitting MS (RRMS), characterized by reversible episodes of neurological deficits and evolve to secondary progressive MS (SPMS) with permanent neurological deficits and continuous progression of disability. Approximately 15% of individuals have a progressive course from disease onset, which is referred to as primary progressive MS (PPMS) ^3^. The precise etiology of MS remains unknown, but epidemiology indicates that it is influenced by both environmental and genetic factors ^3^. Among environmental factors, smoking and low vitamin D are likely contributors ^3,4^, while Epstein-Barr virus infection in early life has been demonstrated to increase risk ^5^. Genetic factors consist of hundreds of small-effect common variants that increase disease risk ^6^ or severity ^7^. Risk variants implicate genes strongly enriched for immune relevance ^6^, while MS severity variants lie in CNS-expressed genes ^7^. This is in line with the well characterized immune-mediated focal losses of myelin in the RRMS phase, in which immune-related therapies readily reduce relapse rates and disease activity. However, efficacy of immune therapies is very limited in the progressive stage characterized by a shift from predominantly localized acute injury to widespread neuroinflammation and neurodegeneration ^8^, apparently originated within the CNS. In fact, the first approved treatment for progressive MS also targets CNS-resident cell types, such as microglia ^9^. Therefore, identifying the CNS-driven molecular determinants of the progressive forms is a current priority in the understanding and management of MS.

A cell type–specific interaction analysis of MS risk loci, used to ascribe pathogenicity of gene variants to cell types, found three loci to be pathogenic in the CNS and specifically in the oligodendrocyte lineage, of which two are involved in promoter-proximal pausing release ^10^. Promoter-proximal pausing is a regulatory mechanism in gene expression where RNAPII temporarily halts transcription shortly after initiation, providing a checkpoint for the regulation of gene expression and facilitating rapid responses to cellular signals ^11^.

TAF1 is the largest subunit of the general transcription factor TFIID, which makes contacts with core promoter DNA elements to aid to define the transcription start site (TSS), and coordinates the formation of the RNAPII transcriptional preinitiation complex (PIC) ^12^. Interestingly, TAF1 participates in RNAPII promoter-proximal pausing ^13^ and its deficiency has been associated to brain disorders like MRXS33 X-linked intellectual disability syndrome ^14^ and X-linked dystonia-parkinsonism (XDP) ^15,16^. Furthermore, we have reported *TAF1* mis-splicing in Huntington’s disease, leading to decreased TAF1 protein stability ^17^. Given TAF1’s role in promoter-proximal pausing ^13^ and neurological disorders ^14–17^, the association of MS severity with variants in CNS-expressed genes ^7^, and the fact that most MS-linked genes that are pathogenic in oligodendrocytes regulate RNAPII pausing ^10^, we decided to analyze TAF1 in brains of progressive MS cases.

## Results

### Decreased detection of the C-terminal end of TAF1 in brains from individuals with MS

We analyzed TAF1 levels in progressive MS brain tissue by Western blot with antibodies against different parts of the protein. Specifically, to determine whether TAF1 is altered where pathogenic progression-driving CNS-resident changes are expected to emerge independently of infiltrating inflammatory cells, we examined cortical samples containing normal-appearing grey and white matter (NAG+WM) from individuals with PPMS or SPMS (table S1). The human *TAF1* gene is located on chromosome X and encompasses 38 exons that encode the 1893–amino acid canonical form of the protein (Uniprot P21675-2) (Fig. 1A). We used three commercial antibodies (Abs) which were raised against the following amino acid sequences: 103 to 123 (encoded by exon 3), 1771 to 1821 (encoded by exon 37), or 1821 to 1871 (encoded by exon 38) (Fig. 1A), which we validated by Western blot on a human cell line transfected with either TAF1 siRNA or human TAF1 over-expressing plasmid (fig. S1). Strikingly, despite unaltered total protein levels of TAF1-as detected with exon3 and exon37 Abs-we found a specifically reduced immunodetection of the C-terminal end of TAF1 with the antibody raised against the sequence encoded by the last exon (ex38 Ab), in both PPMS and SPMS samples (Fig. 1B).

**Fig. 1.**
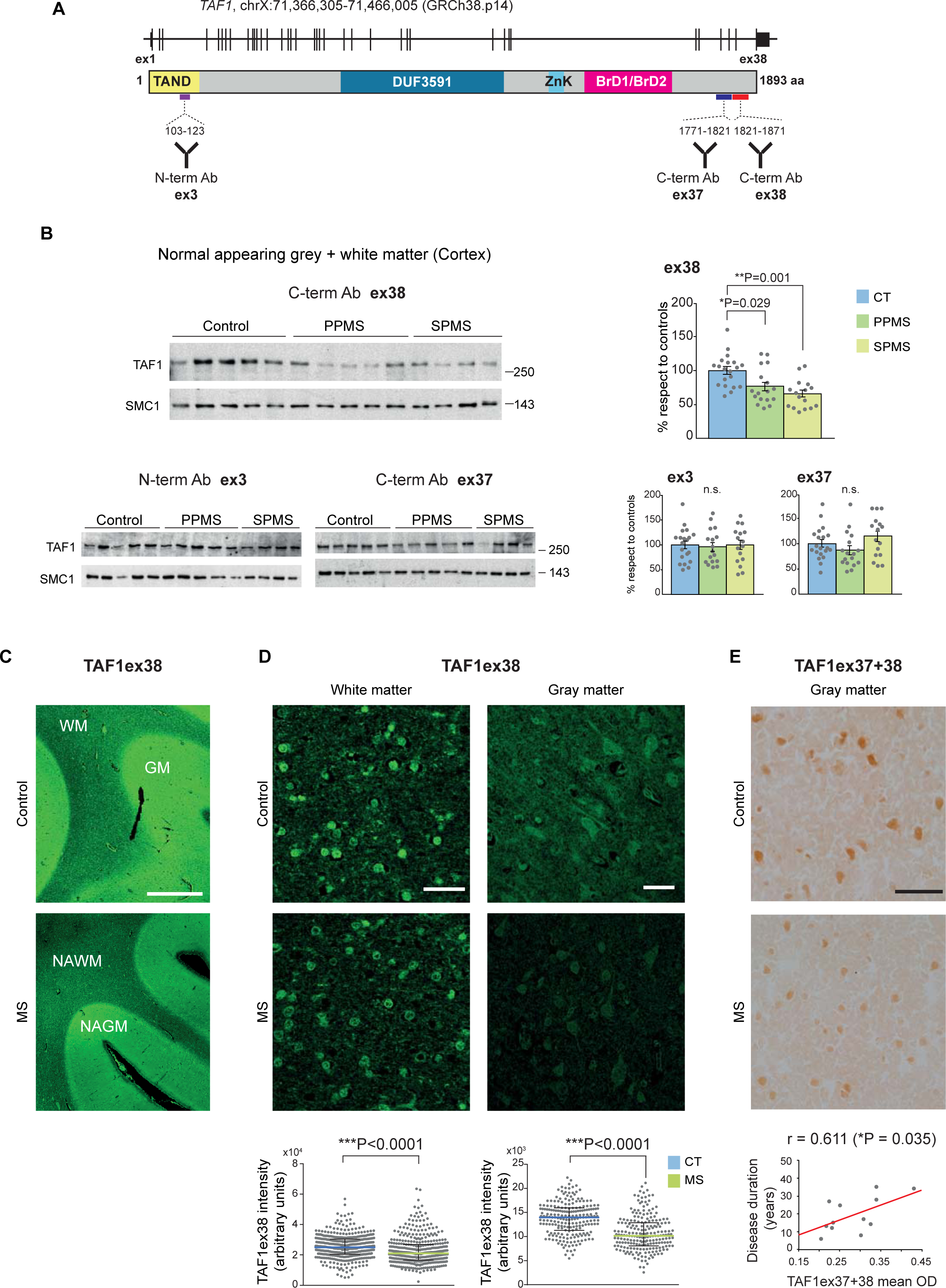
Reduced detection of C-terminal TAF1 in NAG+WM from individuals with MS. (A) Schematic view of human TAF1 gene and protein (Uniprot P21675-2), with the following domains: TAF1 N-terminal domain (TAND), DNA binding domain (DUF3591), zinc knuckle (ZnK) and the bromodomains (BrD1 and BrD2). Also indicated the positions of the protein sequences used as immunogens to generate the antibodies employed in the Western blot analyses, each mapping to a different exon (3, 37 and 38). (B) TAF1 detection with antibodies against sequences encoded by exons 38, 3 or 37 in nuclear cortical extracts (NAG+WM) from controls (CT, n = 20) and individuals with PPMS (n = 17) or SPMS (n = 16), quantified by normalizing respect to SMC1 levels (Kruskal-Wallis test or one-way ANOVA followed by Tukey’s post hoc test). Graphs show mean ± SEM. n.s., non-significant. (C) Immunofluorescence in control and progressive MS cortical sections with TAF1ex38 antibody. Scale bar: 2 mm. (D) TAF1ex38 immunofluorescence in cortical sections from control (n = 6) and MS (n = 6 for WM, 5 for GM) cases and quantification of intensity in 70 and 40 nuclei per case in WM and GM, respectively. Scale bar: 30 µm. Graphs show mean gray intensity quantification in WM/NAWM (left) or GM/NAGM (right) nuclei (median ± interquartile range, Mann–Whitney U test). (E) Immunohistochemistry in control (n = 3) and MS (n = 12) cortical sections with the TAF1ex37+38 antibody. Graph shows simple linear regression (Pearson correlation coefficient, two-tailed significance) with significant correlation between decreased C-terminal TAF1 staining (mean optical density, OD) and lower disease duration of both PPMS and SPMS cases. Scale bar: 50 µm.

To anatomically characterize the decreased biochemical detection of TAF1 with ex38 Ab in NAG+WM, we performed histological analyses on control and MS brain cortical sections with ex38 Ab, as well as the two anti-TAF1 antibodies validated in the Human Protein Atlas, the one raised against the whole protein (TAF1 whole protein Ab) and the one raised against the C-terminal region (ex37+ex38 Ab) (Fig. 1C-E and fig. S2). In control tissue, similar patterns were observed with the three antibodies, which detected TAF1 across most cell types of cerebral cortex, in accordance with data in the Human Protein Atlas. In more detail, most TAF1 positive cells in WM correspond to oligodendrocytes (as evidenced by double labeling with the oligodendroglial marker TPPP), whose staining is essentially restricted to the cell nucleus (Fig. 1D and fig. S2A,B), while neurons in GM showed both nuclear and cytoplasmic staining (Fig. 1D and fig. S2A). The comparison with progressive MS tissue evidenced that TAF1ex38 fluorescence intensity was reduced in both NAGM and NAWM, confirming the decreased signal observed by Western blot, and indicating the contribution of both neurons and oligodendrocytes (Fig. 1C,D). Furthermore, staining with the TAF1ex37+ex38 Ab showed that lower labelling intensity in NAGM significantly correlated with shorter disease duration and, presumably, a more aggressive course (Fig. 1E).

We then performed a bioinformatics analysis of the C-terminal sequence of TAF1. In agreement with a previous prediction using AlphaFold ^18^, the PrDOS tool depicts a disordered region covering the entire sequence after the tandem bromo domains (BrDs) (fig. S3A). We observed a high degree of conservation at the BrDs (encoded by exons 28–32), followed by a valley of low homology. Notably, conservation reappears at the particularly unstructured (disorder probability > 0.6) region downstream of mid-exon 36, suggesting the existence of functional domains at the C-terminal end of TAF1. Finally, the sequence encoded by exon 38 is particularly rich in conserved glutamic and aspartic residues (fig. S3B,C). Together, our biochemical, histological and bioinformatics analyses evidenced decreased detection of the highly conserved, intrinsically disordered and acidic C-terminal end of TAF1 in neurons and oligodendrocytes of normal appearing brain tissue of progressive MS cases.

### Extralysosomal CTSB in MS brains explains decreased detection of C-terminal TAF1

To gain insight into the possible molecular mechanisms behind the decreased detection of C-terminal TAF1 observed in normal appearing brain tissue of progressive MS cases, we performed bioinformatics analyses on TAF1 transcript and protein sequences in search of splicing or proteolysis events able to generate versions of TAF1 protein lacking exon 38-encoded sequences.

Regarding TAF1 transcript variants, there are three reported alternative splicing events (two of which are conserved between mouse and human) that generate isoforms that totally or partially lack ex38 (fig. S4A,B). We performed high depth bulk tissue RNA-seq on NAG+WM from control (n=4), PPMS (n=3) and SPMS (n=3) individuals (fig. S4C) and alternative splicing analyses with three bioinformatic tools (Vast-tools, rMATS and Majiq) failed to detect significant differences on ex38-variants (fig. S4C and table S2). This was confirmed in a larger cohort of control and progressive MS cases through quantitative or digital droplet PCR (fig. S4D). These results indicate that splicing alterations are unlikely to be responsible for the decreased detection of exon38-encoded sequence of TAF1 in MS brain tissue.

Regarding proteolysis, we screened the whole TAF1 protein sequence with the Procleave tool ^19^ for prediction of cleavage sites by 48 different proteases. This revealed a site of cleavage by CTSB with maximal score at the beginning of exon 38-encoded sequence (amino acid 1840, TSFS† SIGG) (table S3). In good agreement, ProsperousPlus ^20^ identified this site, together with two additional CTSB sites, in exon 38-encoded sequence, also with high score (Fig. 2A). CTSB is a lysosomal protease that retains endoproteolytic activity at neutral pH in the cytoplasm upon transient lysosomal permeabilization, a phenomenon often associated to neurodegenerative processes ^21^. Interestingly, increased lysosomal fragility has been reported in MS brain tissue ^22^, and it has also been found that CTSB activity is increased in peripheral blood cells ^23^, cerebrospinal fluid ^24^ and brain tissue ^25^ of MS cases. This prompted us to explore the possible extralysosomal localization of CTSB in normal appearing brain tissue of progressive MS cases by immunofluorescence. As expected, control samples showed a lysosomal-like punctate pattern of CTSB (Fig. 2B). Interestingly, progressive MS samples show a more intense CTSB labeling that extends beyond the lysosomal-like puncta, rendering a diffuse labeling in the cytoplasm and, to a lower extent, the nucleus. Quantification of the latter confirmed increased extralysosomal CTSB signal in both GM and WM of progressive MS cases (Fig. 2B).

**Fig. 2.**
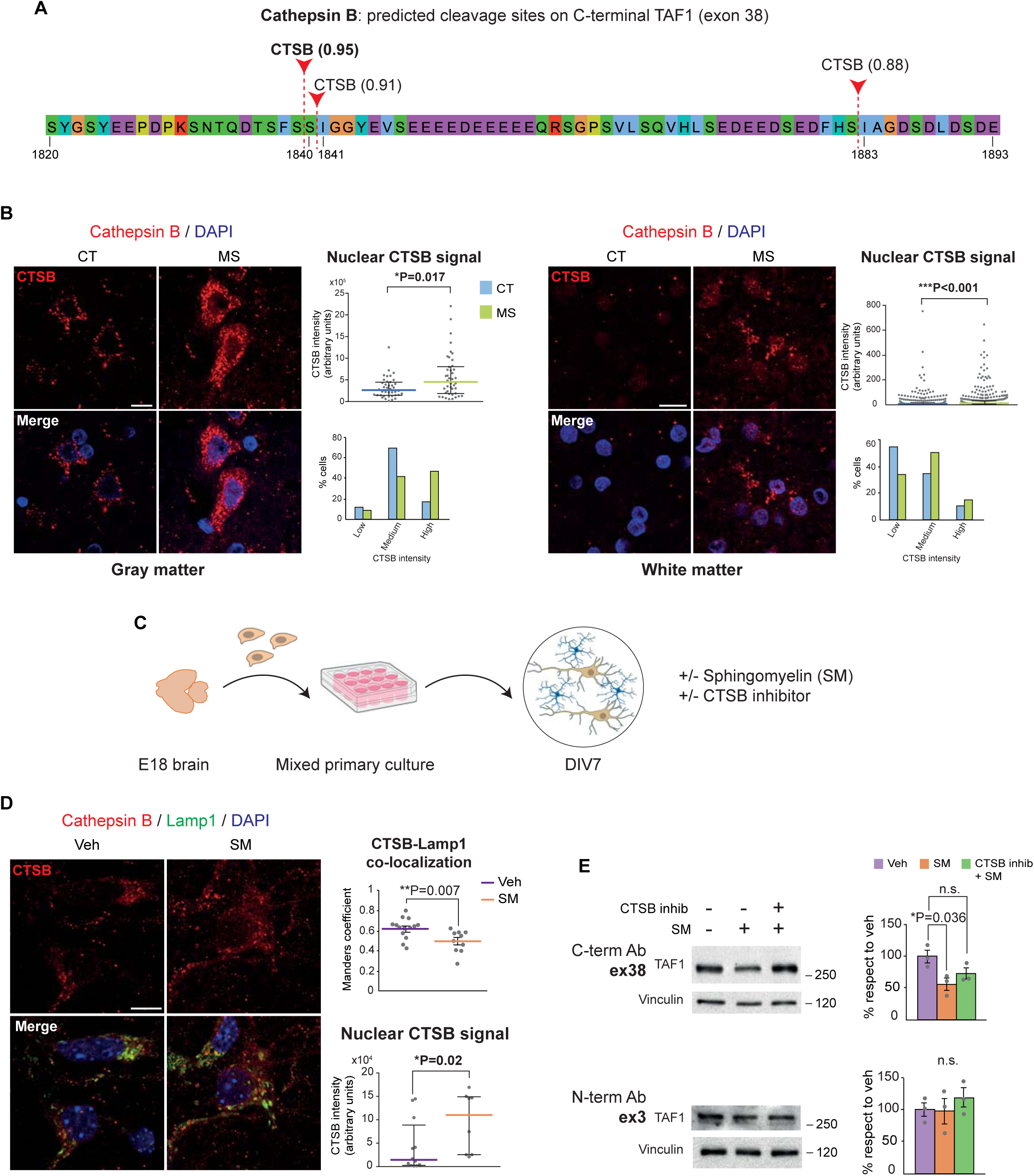
Endoproteolysis by extralysosomal CTSB explains underdetection of C-terminal TAF1. (A) CTSB cleavage sites predicted with ProsperousPlus tool at human TAF1 protein sequence encoded by exon 38 are shown with their respective probability scores in parentheses (range 0-1, threshold 0.8). (B) Immunofluorescence in control (n = 3) and MS (n= 3) cortical sections with CTSB antibody with DAPI nuclear counterstaining, both in gray (left) and white (right) matter. Scale bars: 10 µm. Graphs show quantification of the nuclear CTSB intensity signal (top; median ± interquartile range, Mann–Whitney U test) and its distribution across experimental groups (bottom) in both GM and WM regions. (C) Mixed primary cortical culture from E18 wt mice scheme. Drawings were imported from NIH Bioart. (D) Double immunofluorescence with CTSB and the lysosomal marker Lamp1 with DAPI nuclear counterstaining after treatment with either vehicle (Veh) or 40 µM sphingomyelin (SM). Scale bar: 10 µm. Graphs show Manders’ coefficient quantification for CTSB-Lamp1 colocalization (up; mean ± SEM, two-tailed unpaired t-test) and nuclear CTSB intensity signal (down; median ± interquartile range, Mann–Whitney U test). (E) TAF1 detection with antibodies against sequences encoded by exons 38 or 3 in mixed primary cortical culture extracts after treatment with either vehicle, 40 µM sphingomyelin, or 2 µM of the CTSB inhibitor CA074Me + 40 µM sphingomyelin, quantified by normalizing respect to vinculin levels (one-way ANOVA followed by Tukey’s post hoc test). Graphs show mean ± SEM from 3 independent cultures. n.s., non-significant.

We then investigated in primary neuron/glia mixed cultures whether sphingomyelin-induced transient lysosomal permeabilization and subsequent leakage of CTSB, results in C-terminal modification of TAF1 (Fig. 2C). Sphingomyelin was chosen because the levels of this major component of myelin are increased in periplaque NAWM of MS brains ^26^ and incubation of cell cultures with sphingomyelin is known to induce lysosomal permeabilization and subsequent decreased co-localization of CTSB with the lysosomal marker Lamp1 ^27^. As expected, we observed diminished co-localization of CTSB with Lamp1, together with a spreading of CTSB signal across cytoplasmic and nuclear compartments, similar to that seen in MS tissue (Fig. 2D). Remarkably, this correlates with decreased biochemical detection of TAF1 with the exon 38 antibody, but not with the N-terminal antibody (Fig. 2E). Furthermore, the decreased detection of C-terminal TAF1 was prevented by pretreatment with CA-074Me (a cell permeable prodrug of the CTSB inhibitor CA-074 ^28^) (Fig. 2E). Together, these results point to endoproteolysis by extralysosomal CTSB as a likely mechanism accounting for decreased detection of exon 38-encoded C-terminal end of TAF1 in normal appearing brain tissue from progressive MS cases.

### Progressive disability in mice lacking the C-terminal end of TAF1

To explore the function of the highly conserved sequence at the C-terminal end of TAF1 and its possible pathological relevance *in vivo*, we generated a genetically modified mouse model that mimics the decreased detection of this region in MS brains. For this, we emulated the most prevalent alternative splicing event (19% of *TAF1* transcripts in human cerebral cortex ^29^) that precludes expression of ex38-encoded sequence, namely intron 37 retention which results in truncation of the TAF1 protein right after the exon 37–encoded sequence, with the only change of the last amino acid (S1820R) (fig. S4B). We designed a CRISPR-Cas9 strategy to eliminate the reported splicing acceptor sites in exon38 and in the canonical 3’UTR, obtaining a mouse line with a *Taf1d38* allele in which 4,246 base-pairs were deleted, without introducing any point mutation (Fig. 3A). RNA-seq profiles of hemizygous males or homozygous females for the *Taf1d38* allele showed that the remaining intron 37 sequence and the distal UTR formed a new 3’UTR immediately downstream of the canonical exon 37 (Fig. 3B). By Western blot, we verified that there was no signal with the exon38 antibody, while total TAF1 levels remain unaltered (using the N-terminal [exon3] antibody) (Fig. 3C).

**Fig. 3.**
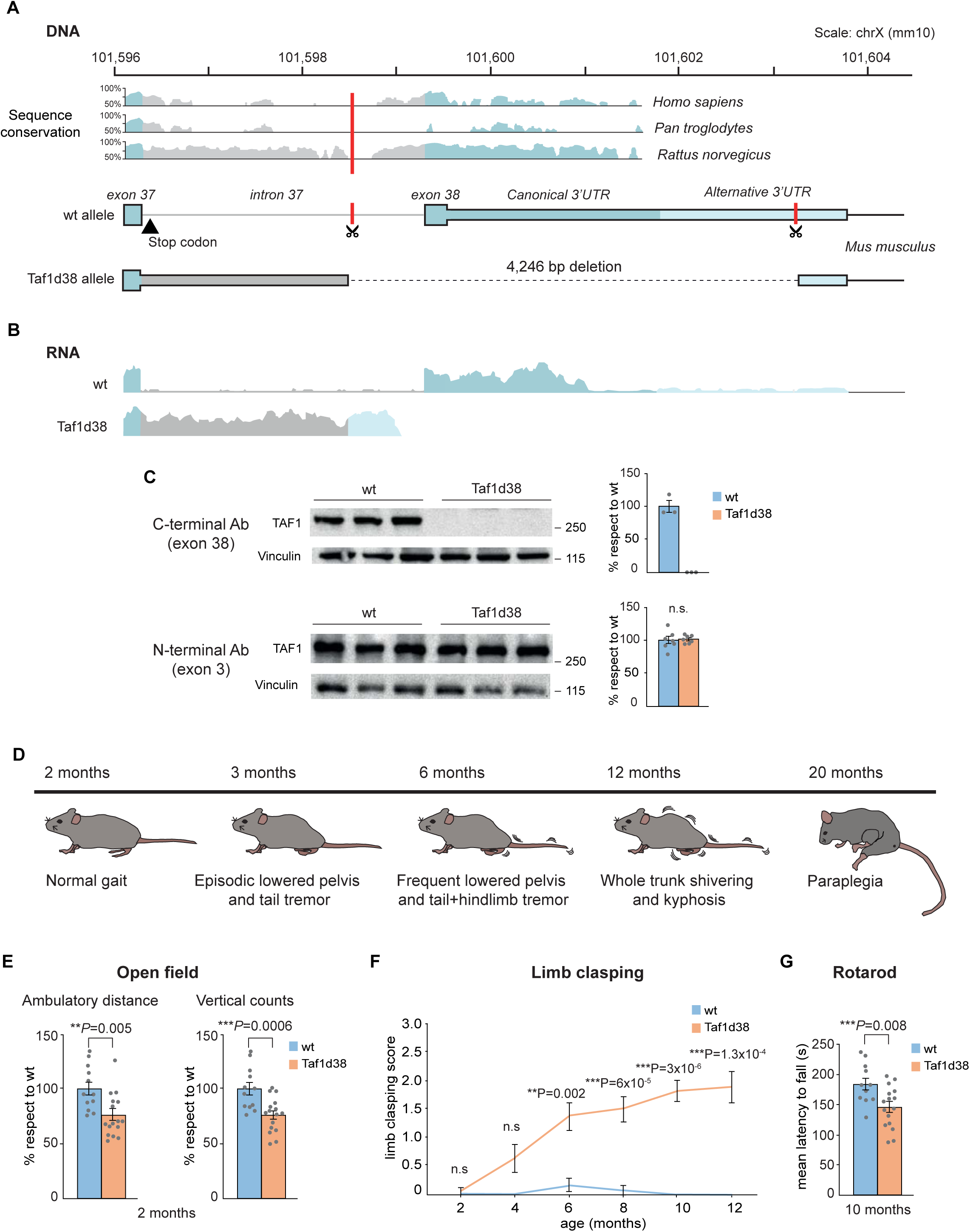
Generation of Taf1d38 mice and behavioral characterization. (A) Schematic view of the 3’ region of TAF1 gene comprising exon 37 to 3’UTR (chromosomal coordinates 101,596-101,604 according to the GRCm38 mm10 assembly). The scissors show the targets of the guide RNA in a region inside intron 37 with low conservation and inside the alternative 3’UTR, leading to a deletion of 4,246 bp. (B) Integrative Genomics Viewer (IGV) profiles of RNA-seq data from a wild-type (wt) and a hemizygous Taf1d38 mouse at the 3’region of TAF1. In the wt mouse, exons 37 and 38 and the canonical 3’UTR (blue) are present, minimal intron 37 retention (gray) is seen and the alternative 3’UTR (light blue) has few reads. In the Taf1d38 mouse, exon 37 (blue) is followed by the initial part of intron 37 (gray) fused to the distal part of the alternative 3’UTR (light blue) which provides a polyA sequence. (C) Detection of TAF1 protein in the brain of 1.5-month-old wt or Taf1d38 mice with the antibodies against sequences encoded by exons 38 (n = 3 for each genotype) or 3 (n = 7 for each genotype). (D) Time course of evident motor phenotype of Taf1d38 mice from early (3 months) to end-stage disease (20 months). (E) Ambulatory distance traveled and number of vertical activity counts in open field test by 2-month-old wt (n = 12) or Taf1d38 mice (n = 16). (F) Limb clasping score evolution from early to symptomatic stages of wt (n = 12 for 2-8 months and n= 11 after 8 months) or Taf1d38 mice (n = 16). Mann-Whitney U test. (G) Mean latency to fall in the rotarod test for 10-month-old wt (n = 11) or Taf1d38 mice (n = 16). (C,E,G) Two-tailed unpaired t-test. (C,E-G) Graphs show mean ± SEM.

Mice carrying at least one *Taf1d38* allele were born at the expected frequencies and they appeared normal until the age of 3 months, when hemizygous males and homozygous females (hereby termed Taf1d38 mice) began to show a visible progressive disability phenotype mainly affecting the hindlimbs, mimicking some of the symptomatology associated with mouse MS models such as the experimental autoimmune encephalomyelitis (EAE) model ^30^. At 3 months, Taf1d38 mice start showing episodes of tail tremor and of lowered pelvis during walking. At 6 months, the action tremor of the tail became permanent and the episodes of lowered pelvis more frequent, with kyphosis starting to be evident. At 10 months, the tail tremor was accompanied by lower body shivering that extended to the whole trunk at 12 months. The phenotype then progressed to paraplegia, which was frequent in 20-month-old mice (Fig. 3D and Movie S1). For this reason, we established 18 months as the humane endpoint for the entire colony of mice with the *Taf1del38* allele (including heterozygote females whose visible phenotype is subtle).

In line with their evident phenotype of progressive disability, Taf1d38 mice also showed decreased activity (both ambulatory and vertical) in the open field test (Fig. 3E), progressive hindlimb clasping (Fig. 3F) and motor coordination deficit in the rotarod test (Fig. 3G). Together, these results indicate that deletion of the C-terminal portion of TAF1, which is underdetected in human MS brains, results in progressive disability in mice.

### Progressive demyelination and WM neuroinflammation in Taf1d38 mice

We then analyzed the status of myelin in the spinal cord of pre-symptomatic (1-month-old) and symptomatic (7-month-old) Taf1d38 mice by transmission electron microscopy. Myelin appearance and G-ratios were normal at one month, while 7-month-old mice show a clear demyelination, as corroborated by the corresponding increased G-ratios in anterolateral, lateral and dorsal columns (Fig. 4A). Remarkably, at 12 months, the myelin deficit in Taf1d38 mice became evident to the naked eye by the marked loss of the whitish appearance of the spinal cord (Fig. 4B), as further corroborated by FluoroMyelin staining (Fig. 4C). Interestingly, demyelination was not restricted to the spinal cord, as in the brain we observed an apparent decreased staining with BlackGold reagent in corpus callosum and superior colliculus at 6 months (Fig. 4D), and a significant decrease of FluoroMyelin staining was observed in corpus callosum and subcortical white matter at 12 months (while not at 3 months) (fig. S5A). Furthermore, electrophysiological recordings of the fornix of 14-month-old Taf1d38 mice showed increased latency selectively for myelinated fibers (fig. S5B).

**Fig. 4.**
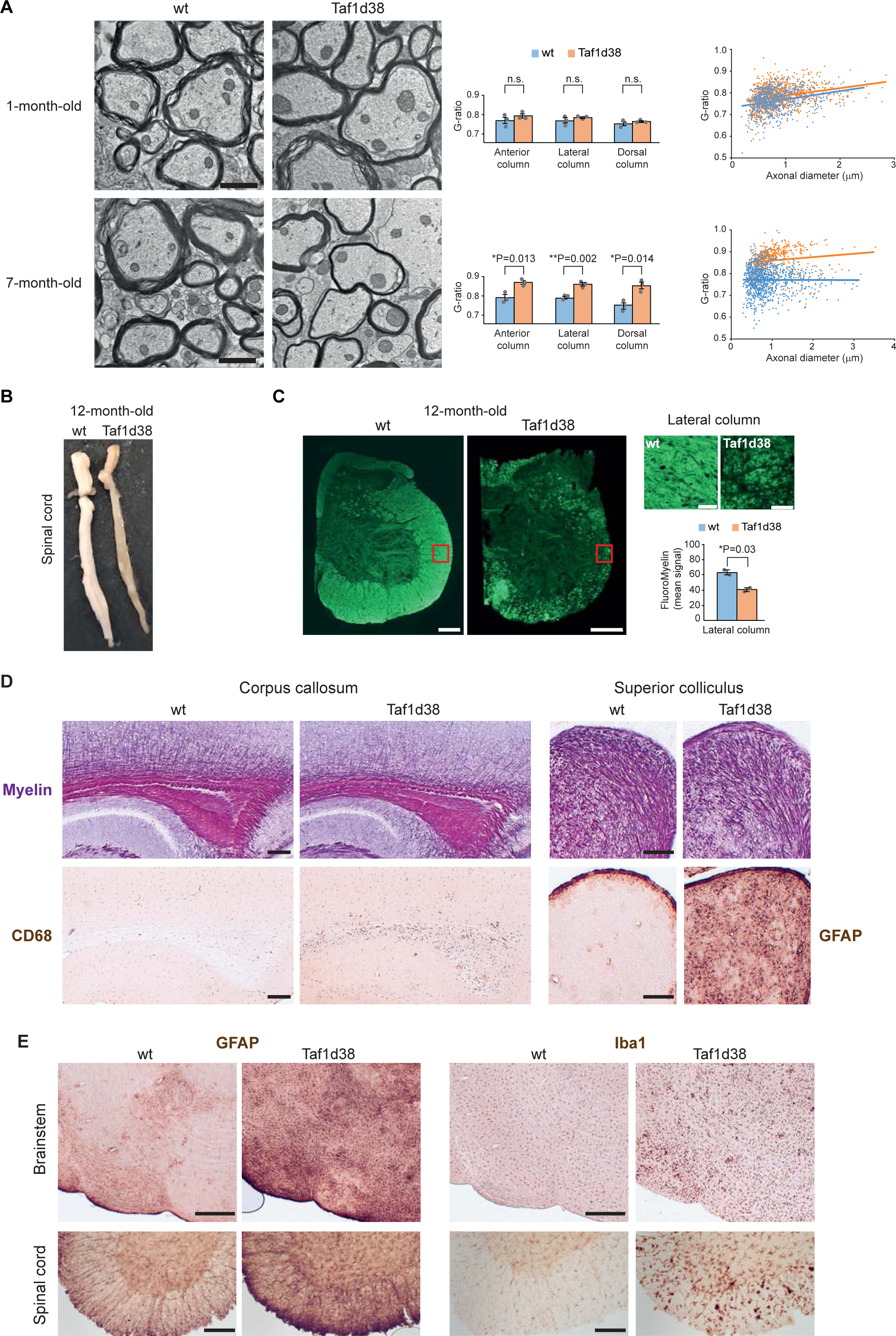
Robust demyelination and WM neuroinflammation in Taf1d38 mice. (A) Transmission electron microscopy of spinal cord WM of 1-month-old (top) and 7-month-old (bottom) mice showing normal myelin formation (top) and posterior marked myelin loss (bottom) in Taf1d38 mice. G-ratio quantification is shown for the three main WM tracts of the spinal cord at thoracolumbar level at both ages (left histogram). Axonal diameter vs G-ratio plot (right) shows that increased G-ratio in 7-month-old Taf1d38 mice occurs independently of axonal size. Twenty microphotographs per tract and per animal (n = 3 for each age and genotype) were analyzed. Scale bar: 1 µm. (B) Photograph of the entire spinal cord of 12-month-old mice showing severe atrophy and WM loss in Taf1d38 mice. (C) FluoroMyelin staining of lumbar spinal cord of 12-month-old mice showing atrophy and profound WM loss in Taf1d38 mice. Insets (boxed areas) in lateral columns are shown enlarged on the right. Histogram shows quantification of the mean signal in the lateral columns of the lumbar spinal cord (n=2). Scale bars: 200 μm (left panels), 10 μm (right panels). (D) Black Gold myelin staining (top) allowing discern myelin loss, and immunohistochemistry staining for CD68 (bottom left) and GFAP (bottom right) showing glial activation, in sagittal sections from 6-month-old Taf1d38 mice. Scale bars: 200 µm. (E) Immunohistochemistry for GFAP (left) and Iba1 (right) in brainstem sagittal sections from 6-month-old (top) and thoracolumbar spinal cord transversal sections from 9-month-old (bottom) Taf1d38 mice, showing marked glial activation. Scale bars: 500 µm (top), 200 µm (bottom). (A,C) Two-tailed unpaired t-test. Graphs show mean ± SD.

To asses neuroinflammation, we performed immunohistochemistry for the microglia/macrophage lineage markers CD68 and Iba1 on brains of 6-month-old Taf1d38 mice. This revealed a profuse staining in most white matter tracts, such as those in the corpus callosum, the superior colliculus, the brainstem and the spinal cord (Fig. 4D,E). A similar pattern was observed after immunostainings for two additional MS neuroinflammation markers, GFAP and Spp1 (osteopontin, OPN) (Fig. 4D,E and fig. S5C). The microglial activation in Taf1d38 mice is not accompanied by infiltration of T or B lymphocytes, as these were not detected by immunofluorescence using the CD3 and B220 markers, not even in the brain regions of Taf1d38 mice with profuse CD68 and Iba1 staining (fig. S5D). Together, these data demonstrate progressive demyelination and CNS resident WM neuroinflammation in Taf1d38 mice.

### MS-like transcriptomic signature in Taf1d38 mice

Transcriptomic analyses were performed to investigate the molecular mechanism underlying the MS-like phenotype of Taf1d38 mice. Since TAF1 is a key subunit of TFIID, we wondered whether removing its C-terminal end would lead to a generalized alteration of transcription or if, on the contrary, it would affect only a minority of genes. We performed bulk tissue RNA-seq of the striatum (a grey and white matter structure that shows atrophy and demyelination in MS ^31,32^) from 2-month-old (before onset of apparent disability) Taf1d38 mice. Strikingly, only 1.8% of genes showed altered transcript levels (Fig. 5A and table S4), with 163 genes downregulated and 131 upregulated. Gene ontology analysis with Ingenuity Pathway Analysis (IPA) on the top differentially expressed genes (DEG) detected *Pathogenesis of MS* and *Vitamin D receptor/retinoid X receptor activation,* as top terms, together with canonical gene sets related to inflammation (Fig. 5B and table S4), thus coinciding with key pathways implicated in MS neuropathology as well as genetic and environmental risk factors ^3,4^. Upregulated genes included markers of neuroinflammation associated to MS, such as *Spp1* (OPN), *Gfap*, *H2-D1*, *H2-K1*, *Ccl4*, *Cxcl10*, *Cd52*, *B2m* and *C4b* (Fig. 5C). In contrast, oligodendroglial-enriched genes ^33^, such as *Ptgds*, *Anln*, *Il33* and *Klk6*, were among the downregulated transcripts (Fig. 5C). Furthermore, the Taf1d38 mouse transcriptomic profile showed a clear overrepresentation of the DEG signature of NAG+WM tissue of individuals with PPMS (Fig. 5D, fig. S4C, 6A,B and table S5), which was particularly remarkable (7-fold enrichment, P < 1.8 × 10^-5^) for the top downregulated genes.

**Fig. 5.**
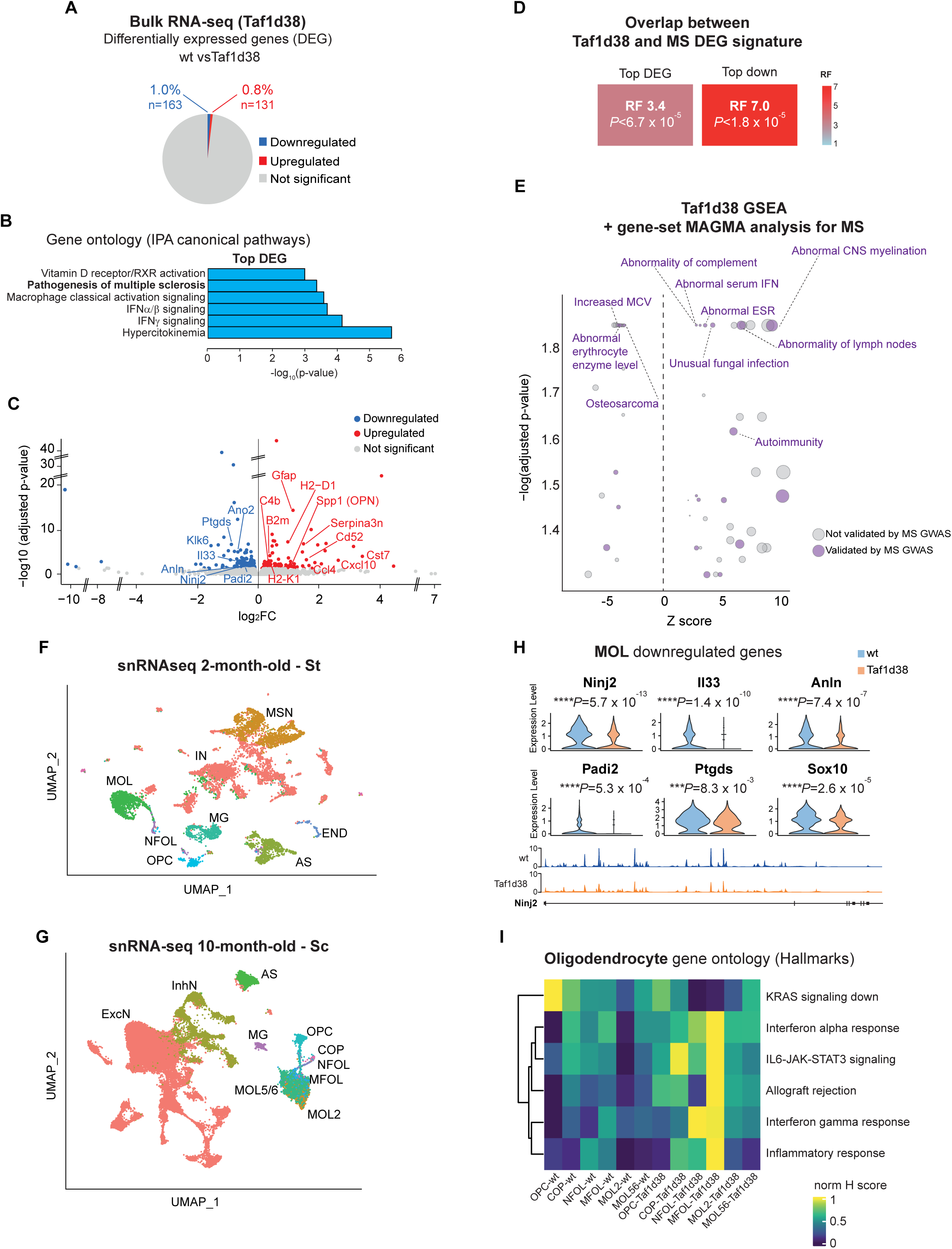
Transcriptomic alterations of Taf1d38 mice. (A) Percentage of DEG identified through bulk tissue RNA-seq analysis of striata from 2-month-old wt (n = 4) or Taf1d38 mice (n = 4). (B) Top significant IPA canonical pathways for the top (log2FC ≥ |0.5|) DEG in Taf1d38 mice. (C) Volcano plot of Taf1d38 DEG. (D) Representation factors (RFs) showing the DEG overlap between Taf1d38 mice and PPMS cortical NAG+WM. The top DEGs were defined for Taf1d38 mice as those with log2FC ≥ |0.25|, and for MS samples as those with log2FC ≥ |1.25|. (E) Bubble plot of top enriched categories from the HPO after GSEA of bulk polyA+ RNA-seq data. The categories that were validated for MS via MAGMA are labeled in purple. ESR, erythrocyte sedimentation rate; MCV, mean corpuscular volume. (F) UMAP plot of cell clusters sorted by cell population after snRNA-seq of the striatum (St) (n = 10,351 nuclei from 3 wt and 3 Taf1d38 mice). (G) UMAP plot of cell clusters sorted by cell population after snRNA-seq of the spinal cord (Sc) (n = 47,649 nuclei from 3 wt and 3 Taf1d38 mice). (H) Top: violin plots of significantly downregulated genes in the MOL population (n = 1,372 nuclei) from striatum. Bottom: IGV track for Ninj2 gene in MOL (merged tracks of 3 replicates). (I) Heatmap of significantly enriched Hallmark categories in Taf1d38 oligodendrocyte populations in spinal cord. (F,G,I) AS, astrocytes; COP, committed oligodendrocyte precursors; END, endothelium; ExcN, excitatory neurons; IN, interneurons; InhN, inhibitory neurons; MFOL, myelin forming oligodendrocytes; MG, microglia; MOL2, mature oligodendrocytes type 2; MOL5/6, mature oligodendrocytes types 5 and 6; MSN, medium spiny neurons; NFOL, newly formed oligodendrocytes; OPC, oligodendrocyte precursor cells.

Using the GWAS summary statistics of the largest meta-analysis thus far (of 47,429 individuals with MS and 68,374 controls) from the International Multiple Sclerosis Genetics Consortium (IMSGC)^6,7^ to perform a gene-based association analysis via MAGMA ^34^, we found that 21.7% of the Taf1d38 DEGs were nominally associated with MS (table S5). We then performed a gene set enrichment analysis (GSEA) of the Taf1d38 DEG dataset against different databases (table S6). In the context of the Human Phenotype Ontology (HPO) database we detected interesting enriched categories and, after validation *in silico* based on MS genetic associations via gene-set MAGMA analysis, we found *abnormal CNS myelination, abnormality of complement* and *abnormal serum IFN* among the top significant ones (Fig. 5E and table S6). Together, these results indicate that C-terminal TAF1 modification selectively affects the expression of a relatively small set of genes, including many potential MS susceptibility genes, resulting in an MS-like transcriptomic signature.

### High transcriptomic perturbation in oligodendroglia of Taf1d38 mice

To investigate the cell types underlying the transcriptional changes unveiled by bulk tissue RNA-seq, we performed single nuclei RNA-seq (snRNA-seq) analyses on CNS regions and time points with low (striatum from 2-month-old mice) and high (spinal cord from 10-month-old mice) degree of affectation. For each tissue, we identified the major cell populations detected in previous snRNA-seq studies ^35,36^ (Fig. 5 F,G; table S7). In young mice, we found that many of the genes detected underexpressed through bulk tissue RNA-seq were downregulated specifically in mature oligodendrocytes (Fig. 5H and table S7). In the highly affected spinal cord samples, we confirmed the downregulation of oligodendroglial genes, such as *Plp1, Hapln2, Sez6l2, Anl*n and *Ptgds* in mature oligodendroglial clusters (table S7), and we found upregulation of inflammatory genes previously associated with MS pathology ^37^, like *Gpnmb, C1qa, C1qb* or *Spp1* across most cell types, including oligodendroglia (fig. S6C and table S7).

To interrogate which cell clusters showed the highest transcriptional disturbance, we performed clustering at higher resolution in both datasets (fig. S6D,E) and found microglial and mature oligodendrocytes subclusters to be the most perturbed in Taf1d38 mice at both ages and tissues (fig. S6F,G and table S7). Finally, we examined the most affected pathways in the oligodendroglial populations at the symptomatic stage through gene set variation analysis (Hallmarks), finding downregulation of *K-Ras signaling* and upregulation of inflammatory related categories such as *interferon-alpha and-gamma response, IL6-JAK-STAT3 signaling* or *inflammatory response*, suggesting the emergence of immune-like phenotypes in Taf1d38 oligodendrocytes (Fig. 5I, table S7). Together, these analyses detected oligodendrocyte populations among those most affected transcriptionally by TAF1 C-terminal modification, while also pointing to microglia affectation.

### C-terminal TAF1 interacts with MS-linked transcription pause release factors

To explore the mechanism by which the lack of TAF1 C-terminal end results in a restricted and MS-like transcriptional alteration, we investigated physiological TAF1 interactors dependent on exon 38-encoded sequence. We performed GFP-Trap immunoprecipitation from an oligodendroglial cell line (Oli-neu) transfected with EGFP-fused-full-length TAF1 (TAF1FL) or-TAF1d38 (fig. S7A), followed by liquid chromatography–mass spectrometry (Fig. 6A and table S8). As expected, most components of TFIID were coimmunoprecipitated (fig. S7B), confirming the validity of our approach at capturing physiological TAF1 interactors. Consistent with the mild transcriptomic alteration in Taf1d38 mice, TFIID components interacted equally with TAF1FL and TAF1d38 (table S8).

**Fig. 6.**
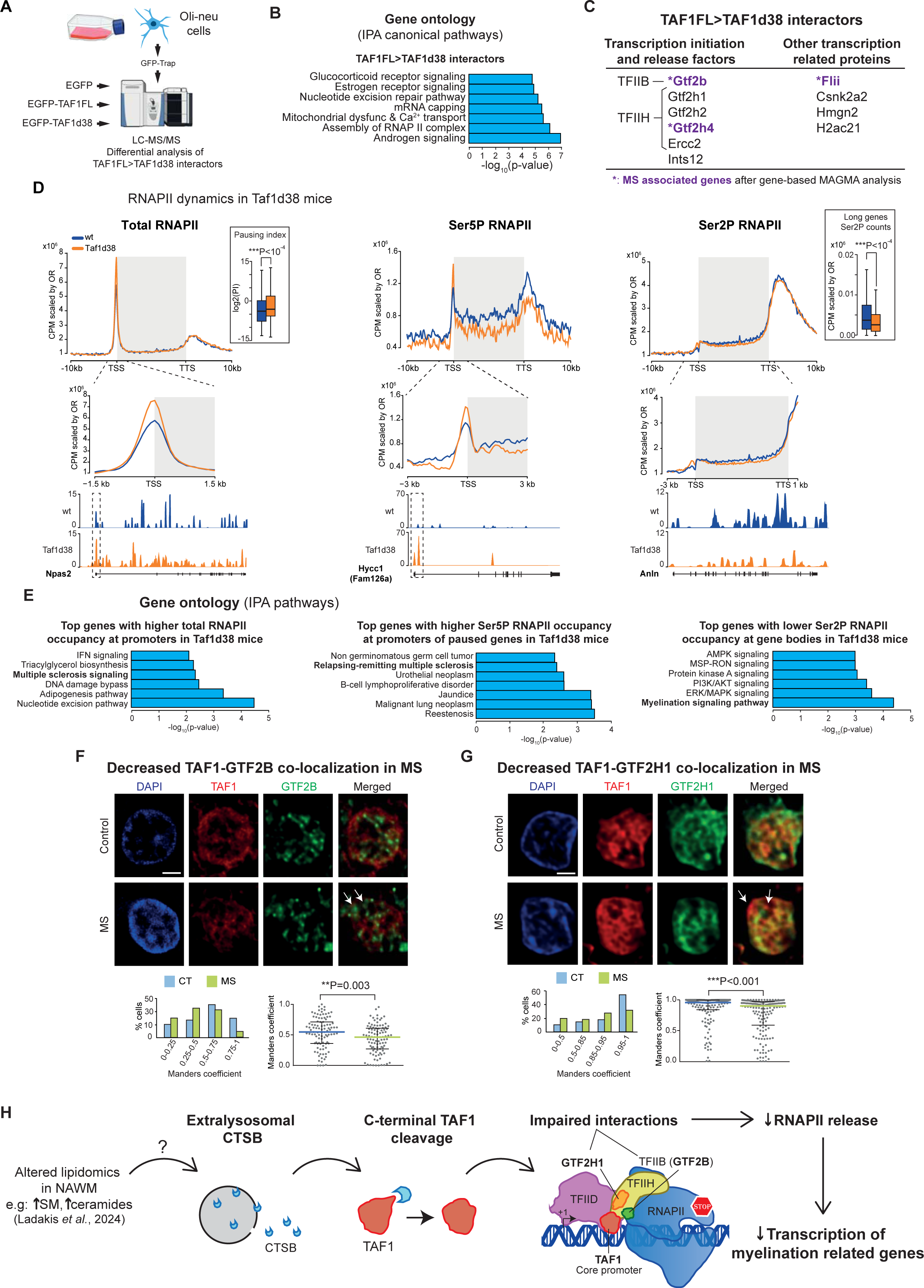
Altered molecular mechanism after C-terminal TAF1 lack. (A) Proteomics experimental design. Oli-neu cells were transfected (in triplicates) with either EGFP, EGFP-TAF1FL or EGFP-TAF1d38 plasmids and processed for GFP-Trap followed by liquid chromatography–mass spectrometry (LC-MS/MS). Drawings were imported from NIH Bioart. (B) Top significant IPA canonical pathways for TAF1FL>TAF1d38 interactors. (C) TAF1FL>TAF1d38 interactors with transcriptional functions. Shown in purple those associated to MS, after gene-based analysis by MAGMA. (D) Total RNAPII (left), Ser5P RNAPII (middle) and Ser2P RNAPII (right, genes larger than 30 kb) signal (spike-in normalized CPM values, scaled by OR) after bulk ChIP-seq of 2-month-old mice (n =2). Focus on the promoter (left and middle) or the gene body (right) region is shown below. The box plot shows the distribution of log2(pausing index) (left) and Ser2P RNAPII normalized RPGCs on gene bodies (right). Bottom: normalized BigWig track of total RNAPII signal along Npas2 (left), Ser5P RNAPII signal along Hycc1 (Fam126a) (middle), and Ser2P RNAPII signal along Anln (right) genes. (E) Top significant IPA pathways for genes whose promoter occupancy by total RNAPII is higher in Taf1d38 mice (left), genes with increased pausing indexes whose promoter occupancy by Ser5P RNAPII is higher in Taf1d38 mice (middle), and genes whose gene body occupancy by Ser2P RNAPII is lower in Taf1d38 mice (right). (F) Double immunofluorescence with TAF1 and GTF2B with DAPI nuclear counterstaining in control (n = 3) and MS (n = 7) WM/NAWM nuclei show decreased co-localization in MS (arrows). (G) Double immunofluorescence with TAF1 and GTF2H1 with DAPI nuclear counterstaining in control (n = 3) and MS (n = 3) WM/NAWM nuclei show decreased co-localization in MS (arrows). (H) Proposed molecular mechanism for progressive MS. (D,F,G) Mann–Whitney U test. Box plots and graphs show median±interquartile range. (F,G) Graphs show Manders’ coefficient distribution across experimental groups (left) and quantification (right). Scale bar: 2 µm.

Regarding interactors that bound more efficiently to TAF1FL than to TAF1d38 (TAF1FL>TAF1d38 interactors), gene ontology analysis detected *assembly of RNAPII complex* among top significant terms, as well as *nuclear receptor (androgen-, estrogen-and glucocorticoid-) signaling* (Fig. 6B and table S8). Among these differential interactors, we noticed TFIIB (Gtf2b) and multiple components of the TFIIH complex [i.e. Gtf2h1 (p62), Gtf2h2 (p44), Gtf2h4 (p52) and Ercc2 (XPD, p80)]. Interestingly, both TFIIB and TFIIH are part of the PIC and participate in regulating RNAPII promoter-proximal transcription ^38–40^ (Fig. 6C). Besides, Inst12, which is a subunit of the Integrator complex that regulates transcriptional initiation and pause release following activation ^41,42^ was also detected as a TAF1FL>TAF1d38 interactor, together with other proteins involved in transcriptional regulation such as Flii, Hmgn2, H2ac21 and Csnk2a2 (Fig. 6C).

The above mentioned data indicate that the C-terminal end of TAF1 interplays with multiple transcriptional regulators involved in RNAPII transcriptional initiation and pause release, a mechanism that has been genetically associated to MS specifically in oligodendrocytes ^10^, as mentioned. In this regard, upon gene-based MAGMA analysis of the GWAS data from the IMSGC ^6,7^, we found that the detected TAF1FL>TAF1d38 interactors Gtf2h4 and Gtf2b showed genetic nominal association with MS risk and severity, respectively (Fig. 6C and table S8); the same applies to Flii (Flightless), an actin remodeling protein and transcriptional coactivator of hormone-activated nuclear receptors ^43^, for MS risk.

### RNAPII promoter-proximal pausing at MS-related genes

To characterize molecular details governing the transcriptional dysregulation associated to the loss of TAF1 C-terminal end, we analyzed the distribution of total, initiating (Ser5P) and elongating (Ser2P) RNAPII in brain tissue of 2-month-old Taf1d38 or wild-type mice by ChIP-seq. Globally, we observed an increase of total RNAPII specifically at promoter regions (Fig. 6D), but an unaltered occupancy profile along gene bodies and transcription termination sites (TTS), resulting in increased RNAPII pausing index (Fig. 6D and table S9). In good agreement, we also found an increase in Ser5P RNAPII signal at promoters (Fig. 6D). Gene ontology analysis of the genes displaying a higher increase in total or Ser5P RNAPII levels at the promoter regions showed *Multiple sclerosis signaling* and *Relapsing-remitting multiple sclerosis* among top affected pathways (Fig. 6E and table S9). In line with the increase in RNAPII promoter-proximal pausing, we observed decreased Ser2P RNAPII levels along gene bodies (Fig. 6D). Gene ontology analysis of top genes with decreased Ser2P RNAPII gene body occupancy detected *Myelination signaling pathway* as the most significant terminal (Fig. 6E and table S9). These changes in RNAPII dynamics were observed in the four quartiles of total gene body RNAPII or Ser2P RNAPII occupancy (fig. S7C,D), suggesting global trends, rather than effects on selective sets of genes. Together these data suggest that defects in promoter escape or promoter-proximal pause release take place in the Taf1d38 mouse cells, markedly affecting myelination-and MS-related genes.

### Decreased TAF1-GTF2B and-GTF2H1 co-localization in MS brains

Upon noticing that TFIIB and TFIIH subunits that display exon-38 dependent TAF1 interaction show genetic association with MS risk and severity, and the altered RNAPII dynamics at myelination-related genes upon C-terminal TAF1 modification *in vivo*, we wondered whether the TAF1 C-terminal end defect in MS brains would correlate with diminished co-localization of TAF1 with TFIIB and TFIIH subunits in oligodendrocytes. In fact, by double immunofluorescence on sections from control and progressive MS cases we confirmed significantly diminished signal co-localization for both GTF2B (Fig. 6F) and GTF2H1 (Fig. 6G) with TAF1 in NAWM nuclei of PPMS and SPMS cases. In summary, these proteomic, ChIP-seq and immunofluorescence data provide mechanistic insight evidencing that the C-terminal alteration of TAF1 observed in brain tissue from individuals with MS synergizes with transcriptional initiation and pause release factors genetically associated to MS, to cause decreased expression of oligodendroglial myelination–related genes.

## Discussion

The central finding in this study is the underdetection of the C-terminal end of TAF1 in postmortem brains of individuals with progressive MS (Fig. 6H). Besides, we provide evidence that this may be attributable to endoproteolytic cleavage by CTSB, as we observed increased extralysosomal CTSB in MS brains and demonstrate that induction of lysosomal permeabilization with sphingomyelin suffices to cause CTSB-dependent decrease of the C-terminal end of TAF1 (Fig. 6H). Interestingly, sphingomyelin is a major component of myelin and its levels and those of ceramides are increased in periplaque NAWM of MS brains ^26^, probably as a consequence of myelin breakdown, thus providing a possible mechanism for spreading C-terminal TAF1 alteration which, in turn, leads to decreased interaction with transcriptional initiation and pause release factors genetically linked to MS. Eventually, this results in increased RNAPII promoter-proximal pausing, that markedly affects oligodendroglial myelination–related genes (Fig. 6H).

Besides, by generating the Taf1d38 mouse model, we *in vivo* demonstrate that such C-terminal end alteration of TAF1 suffices to induce a restrictive MS-like brain transcriptomic signature alongside CNS-resident inflammation and young adult-onset progressive demyelination and disability. Compared to other TAF1-modified animal models such as knock-out zebrafish ^44^ or mice ^45^ that show embryonic lethality (see ^46^ for an exhaustive revision), Taf1d38 mice develop normally to adulthood, probably because the absence of the C-terminal end of TAF1 results in a very restricted transcriptomic alteration. Thus, Taf1d38 mice show construct and face validity for modeling progressive MS, as they incorporate the biochemical alteration found in MS brains, and exhibit spontaneous disease onset in young adults, together with genuine progression. This contrasts with other MS models such as EAE, where chronic stages merely reflect the stabilization of severe neurological deficits rather than true worsening over time^47^.

Regarding other known TAF1-related diseases, XDP is characterized by decreased levels of the full protein due to intron 32 retention caused by the retrotransposon insertion ^15^. However, this also results in diminished usage of exons downstream of intron 32 ^15^, such as the 34’ microexon ^16^ - although the latter has more recently been ruled out through long-read sequencing of *TAF1* mRNAs from XDP and control brains tissues ^48^. In any case, given the reduced expression levels of TAF1 exons downstream of the retrotransposon it would be interesting to explore a potential imbalance in detection with N-and C-terminal TAF1 antibodies in XDP brain tissue, similar to the one report here regarding MS. It is worth noting that XDP neuropathology, apart from the loss of striatal medium spiny neurons ^49^, also includes overall changes of the structural integrity of white matter ^50^.

By identifying TAF1 to be a key player in MS pathogenesis, this study has important etiological and translational implications for MS and provides novel therapeutic targets. TAF1 itself is a clear candidate, but we also provide mechanistic clues upstream and downstream of TAF1 modification that point to additional possible targets, like the extralysosomal CTSB and the pausing-release machinery that seems to be particularly compromised in oligodendroglia. In this sense, lysosomal membrane stabilizers like arimoclomol ^51^, CTSB inhibitors such as VBY-376 (whose safety has already been tested in a clinical trial for hepatic fibrosis, https://clinicaltrials.gov/study/NCT00557583) ^52,53^, or even potential competitive peptide inhibitors targeting the CTSB cleavage site in C-terminal TAF1, might be beneficial. Remarkably, the Taf1d38 genetic mouse model of MS represents a useful tool to preclinically test strategies with potential to alleviate transcriptional pausing of myelination related genes, or any other intervention likely to prevent or halt the progressive phase of the disease.

In summary, this study provides, to the best of our knowledge, the first reported alteration of the transcription machinery in the brain of individuals with MS that, when placed into an animal model, suffices to induce a full, progressive MS-like phenotype. Therefore, impaired interactions between the C-terminal end of TAF1 and transcriptional initiation and pause-release factors genetically linked to MS - with subsequent reduced expression of oligodendroglial myelination genes-, emerge as an unprecedent molecular mechanism in MS, which offers multiple novel avenues for exploring therapeutic opportunities.

## Materials and Methods

### Human brain tissue samples

Human brain samples were provided by the UK Multiple Sclerosis Tissue Bank at Imperial College (London) and the Netherlands Brain Bank at the Netherlands Institute for Neuroscience (Amsterdam). Written informed consent was obtained from brain donors and/or next of kin for brain autopsy and for the use of material and clinical data for research purposes. Clinical and neuropathological diagnosis of MS was confirmed for all patients, and the clinical course was defined as secondary or primary progressive by a certified neurologist according to McDonald or Poser criteria. Controls were chosen among those available to best match age, sex and postmortem interval with MS cases, avoiding those with diagnosis of any other neurodegenerative disease. Donor metadata are summarized in table S1. The current histological, biochemical and molecular study with human samples has been approved by the CSIC Ethics Committee (Madrid, Spain).

### Animals

CRISPR mice lacking exon 38 of Taf1 (Taf1d38) and FL_TAF1 mice were generated for this study and used in a C57BL/6J background. All the experiments with Taf1d38 mice were performed between the 5^th^ and the 11^th^ backcross to C57Bl/6j background. The phenotype has remained unaltered to the current 13^th^ backcross. All mice were housed in the CBM animal facility under *ad libitum* food and water availability, and maintained in a temperature-controlled environment on 12 h light–dark cycles, following local authority guidelines approved by the CBM Institutional Animal Care and Utilization Committee (Comité de Ética de Experimentación Animal del CBM, CEEA-CBM), and Comunidad de Madrid PROEX 293/15 and PROEX 247.1/20. Mice were euthanized with CO2.

### Generation and genotyping of Taf1d38 mice

Taf1d38 mice were generated in a 50% C57BL/6J and 50% CBA/CA background via CRISPR/Cas9-mediated gene editing, by the CNB-CBM mouse transgenesis facility. We mimicked the retention of intron 37 (https://vastdb.crg.eu/event/HsaINT1043632@hg38), which results in a TAF1 protein truncated at the end of exon 37 (fig. S4A,B). We designed a CRISPR-Cas9 strategy to eliminate the reported splicing acceptor sites at exon 38 and the canonical 3’UTR, using a guide RNA mapping to intron 37 (5’: CCAAGTTGTGAAGGACGTAAATG) and another mapping downstream of the canonical 3’UTR (3’: GGGCTACAAGCCCCTAACTTAGG), the latter within an adjacent and longer cryptic alternative 3’UTR (Fig. 3A). The gRNAs were co-injected into fertilized oocytes with Cas9 protein. Confirmation of the correct 4,246 bp deletion without introducing any point mutations was performed through Sanger sequencing.

RNA-seq evidenced the clean deletion and the constitution of a new 3’UTR downstream the canonical exon 37 (Fig. 3B). Taf1d38 mice were genotyped using primers that anneal in intron 37 before the deletion and in the distal 3’ UTR after the deletion (GGCAATTCAGTCTTTGTGTATAGG; AGCTCTAGCTGACCTTAAACTCC).

### Bioinformatic analysis of C-terminal TAF1

Genomic and protein sequences used in the alignments were downloaded from <u>Ensembl</u> and <u>UniProt</u> public databases. Sequence alignments were done with state-of-the-art aligner ClustalO ^54^. JalView 2.11.0 ^55^ was used for sequence alignment editing, visualization and analysis. Chosen conservation measure for protein, among JalView’s repertoire, was Valdar’s conservation ^56^. Protein DisOrder prediction System (PrDOS ^57^) was used to study the intrinsic disorder of TAF1. Procleave ^19^ and its update version ProsperousPlus ^20^ was run to find putative protease cleavage sites in TAF1. Experimentally determined post-translational modifications were searched through PhosphoSitePlus ^58^, and predicted phosphorylation sites were found with NetPhos ^59^ and ELM ^60^.

### Western Blot

Mouse brains were quickly dissected on an ice-cold plate and the different structures stored at-80 °C. Extracts were prepared as described ^61,62^ by homogenizing the brain areas in ice-cold Extraction Buffer (20 mM HEPES pH 7.4, 100 mM NaCl, 5 mM EDTA, 20 mM NaF, 1% Triton X-100, 1 mM sodium orthovanadate, 1 μM okadaic acid, 5 mM sodium pyrophosphate, 30 mM β-glycerophosphate and protease inhibitors [Complete, Roche]). Homogenates were centrifuged at 15,000g for 15 min at 4 °C, and supernatant protein content was determined by Quick Start Bradford kit assay (Bio-Rad, 500-0203).

Protein extracts from 293T cells or primary mixed cultures were prepared by scrapping cells in ice-cold Extraction Buffer.

Samples from human brain were stored at-80 °C and ground with a mortar in a frozen environment resulting in tissue powder. Nuclear extracts from human tissue were prepared by homogenizing the tissue powder in ice-cold extraction buffer A (10 mM HEPES pH 7.9, 10 mM KCl, 0.1 mM EDTA, 0.1 mM EGTA, 2.5 mM DTT, 1 mM sodium orthovanadate, 1 μM okadaic acid, 5 mM sodium pyrophosphate, 30 mM β-glycerophosphate and protease inhibitors [Complete, Roche]). After adding 1% NP40 and incubating in ice for 5 min, the samples were centrifuged at 4,000g for 1 min at 4 °C. The pellet was washed in extraction buffer A and resuspended in ice-cold extraction buffer C (20 mM HEPES pH 7.9, 25% glycerol, 400 mM NaCl, 1 mM EDTA, 1 mM EGTA, 2.5 mM DTT, 1 mM sodium orthovanadate, 1 μM okadaic acid, 5 mM sodium pyrophosphate, 30 mM β-glycerophosphate and protease inhibitors [Complete, Roche]). After sonication (two pulses of 10 s and 100% amplitude, Labsonic M, Sartorius), 1% NP40 was added, and after an incubation of 5 min in ice, the extracts were centrifuged at 15,000g for 5 min at 4 °C. The supernatant constituted the nuclear extract, and protein content was determined by Quick Start Bradford kit assay (Bio-Rad, 500-0203).

Between 10 and 30 μg of protein was electrophoresed on 6% SDS–polyacrylamide gel, transferred to a nitrocellulose blotting membrane (Amersham Protran 0.45 μm, GE Healthcare Life Sciences, 10600002) and blocked in TBS-T (150 mM NaCl, 20 mM Tris–HCl, pH 7.5, 0.1% Tween 20) supplemented with 5% non-fat dry milk. Membranes were incubated overnight at 4 °C with either rabbit anti-TAF1ex3 (1:500; Abcam, ab188427), rabbit anti-TAF1ex37 (1:500; Bethyl, A303-504A), rabbit anti-TAF1ex38 (1:500; Bethyl, A303-505A), rabbit anti-SMC1 (1:2,000; Bethyl, A300-055A), rabbit anti-vinculin (1:15,000; Abcam, ab129002) or mouse anti-β actin (1:50,000; Sigma, A2228) in TBS-T supplemented with 5% non-fat dry milk. After washings with TBS-T, the membranes were incubated with secondary HRP-conjugated anti-rabbit IgG (1:2,000, DAKO, P0448) or anti-mouse IgG (1:2,000, DAKO, P0447), and developed using the ECL detection kit (Perkin Elmer, NEL105001EA). Images were scanned with densitometer (Bio-Rad, GS-900) and quantified with ImageJ 1.52d software.

### Immunohistochemical and myelin staining analyses *Human tissue*

Postmortem control and MS cortical snap-frozen or paraffin-embedded tissue was analysed (table S1). Tissue sections were stained for proteolipid protein (PLP) and human leucocyte antigen (HLA-DR/DQ, referred to as HLA), and MS lesions were classified according to demyelination and cellular distribution as described ^63–66^.

### Immunohistochemistry of snap-frozen tissue

The immunohistochemistry protocol was carried out as previously described ^64^. Briefly, cryosections (10µm, Leica) were thawed and fixed in 4% paraformaldehyde (PFA). After antigen retrieval with citrate buffer 0.1M pH 6 at 90 °C, endogenous peroxidase was quenched with 10% methanol and 3% H2O_2_ and the slides were blocked (5% normal goat serum [NDS], 0.2 % Triton X-100 in PBS). Slides were incubated with rabbit anti-TAF1ex37+38 (1:100; Sigma, HPA001075) overnight. Anti-rabbit biotinylated (1:200; Vector Labs) secondary antibody was used, followed by amplification with Vectastain Elite ABC reagent (Vector Labs) and developing with 0.05% 3,3‘-diaminobenzidine (DAB, Sigma-Aldrich) and 0.003% H_2_O_2_. Images were captured on a BX61 microscope (Olympus) equipped with a color camera CX9000 (MBF). For quantification of TAF1 ex37+38 intensity, a mean of 5 ROIs/sample in GM regions defined throughout all the cortical layers (1.78 ± 0.18 mm^2^/ROI) were used. TAF1 intensity (mean value of the optical density obtained from all TAF1^+^ cells/sample) was measured using QuPath software.

### Immunohistochemistry of paraffin-embedded tissue

After deparaffinisation and rehydration through sequential immersions in xylene and ethanol, sections were blocked for endogenous peroxidase using 0.3% H_2_O_2_ in PBS. Antigen retrieval was performed by heating sections submerged in 10 mM sodium citrate pH 6 to 120°C in an autoclave. Primary antibodies [rabbit anti-TAF1ex38 (1:100, Bethyl, A303-505A), mouse anti-TAF1 whole protein (1:50, SantaCruz, sc735)] were incubated overnight at room temperature (RT), detected using EnVision (Agilent Dako, Glostrup) and developed with DAB (Agilent Dako), and counterstained with haematoxylin. Images were obtained by scanning the slides using an Olympus VS200 system.

### Immunofluorescence of paraffin-embedded tissue

The sections were processed as described for immunohistochemistry, blocked (2% NDS, 0.1% Triton X-100, 4% BSA in PB) and incubated overnight with rabbit anti-TAF1ex38 (1:100, Bethyl, A303-505A), mouse anti-TAF1 whole protein (1:100, SantaCruz, sc735), rabbit anti-TAF1ex3 (1:200; Abcam, ab188427), rabbit anti-GTF2B (1:400; Sigma, HPA061626), mouse-antiGTF2H1 (1:100; Abnova, H00002965), rabbit anti-TPPP (1:1,600, Abcam, AB92305) and rabbit anti-CTSB (1:100; Sigma, HPA048998). Primary antibodies were detected using either EnVision (Agilent Dako, Glostrup) or an HRP conjugated secondary antibody (Southern Biotech) followed by tyramide amplification (1:50 or 1:100, Akoya). For double immunofluorescences, the samples were treated with 0.1 M glycine pH 2, blocked and incubated with other primary antibody, which was detected using either an HRP conjugated secondary antibody followed by tyramide amplification, or an Alexa conjugated antibody (Invitrogen). Autofluorescence was dampened with 0.1% Sudan Black B in 70% ethanol, and the sections were counterstained with DAPI. For CTSB stainings, permeabilization was carried with 0.1% saponin instead of Triton X-100. Images were acquired on a Zeiss LSM710 confocal vertical system. For quantification of the signal intensity, mean gray value and integrated density were measured with Fiji, as previously described ^67^. For colocalization analysis, the images were deconvoluted with Huygens 24.10 (SVI) software suite, and the Mander’s coefficient was determined using the JACoP plugin (Fiji).

### Mouse tissue

Mice were deeply anaesthetized with intraperitoneal Dolethal (2 mg/g) and perfused transcardially with NaCl followed by 4% PFA. Brains and spinal cords were fixed overnight in 4% PFA, cryopreserved for 72 h in 30% sucrose in PBS, added to optimum cutting temperature compound (Tissue-Tek, Sakura Finetek Europe), frozen and stored at-80 °C until use. 30 μm sagittal (brain) or 20 μm transversal (spinal cord) sections were cut on a cryostat (Thermo-Fisher Scientific), collected and stored free floating at-20 °C in 30% glycerol, 30% ethylene glycol in 0.02 M PB.

### Immunohistochemistry

Endogenous peroxidase was quenched with 0.1% H_2_O_2_ in PBS and the sections were blocked (0.5% fetal bovine serum [FBS], 0.3% Triton X-100, 1% BSA in PBS) and incubated with either rat anti-CD68 (1:200; Abcam ab53444), rabbit anti-Iba1 (1:1,000; Wako 019-19741) or rabbit anti-GFAP (1:20,000; Novus NB300-141) overnight at 4 °C. Sections were incubated with biotinylated anti-rat or anti-rabbit secondary antibody, amplified with Vectastain Elite ABC (Vector Laboratories) and developed with DAB (Sigma-Aldrich, D4293). Images were captured with a DMC6200 CMOS camera.

### Immunofluorescence

Sections were blocked (4% NDS, 0.1% Triton X-100 in PB) and incubated overnight at 4 °C with either mouse anti-SPP1 (1:150; Biotechne MAB808), rat anti-CD3 (1:50; BioRad MCA1477) or rat-anti-B220 (1:200; BD Pharmingen clone RA3-6B2), followed by incubation with a secondary fluorochrome-conjugated antibody (Thermo-Fisher Scientific) and Hoechst 33258 (1.5 µg/mL) counterstaining. A Leica Stellaris 5 confocal microscope was used to capture the images.

### FluoroMyelin

Brain or spinal cord sections were mounted and let dry. After hydration in PBS, staining in FluoroMyelin™ (Green Fluorescent Myelin Stain F34651, Molecular Probes) was performed according to the manufacturer’s protocol. Images were captured in a fluorescence microscope (Axiovision, Zeiss) and ImageJ software was used to measure the mean intensity gray value in each region, analyzing one level per animal.

*Black gold.* Mouse sections were mounted and let dry overnight. After rehydration in water, the slides were incubated with Black-Gold II reagent at 65 °C, and fixed with sodium thiosulfate solution according to manufacturer’s instructions (Black Gold II Myelin Staining Kit, Avantor BSENTR-100-BG). A DMC6200 CMOS camera was used to capture the images.

### Transmission electron microscopy

1-month-old and 7-month-old mice (3 wt and 3 Taf1d38 for each age) were deeply anaesthetized with intraperitoneal Dolethal (2 mg/g) and perfused transcardially with NaCl followed by 4% PFA and 2.5% glutaraldehyde in PB. Spinal cords were dissected and 2 mm long lumbar fragments were postfixed overnight at 4 °C. Fragments were immersed in 1% osmium tetroxide in 0.1 M cacodylate buffer, dehydrated in ethanol and embedded in Epon-Araldite. Ultrathin transversal sections were collected on pioloform-coated grids and stained with uranyl acetate and lead citrate. The sections were analyzed with a JEM-1010 transmission electron microscope (Jeol, Japan) equipped with a side-mounted CCD camera Mega View III (Olympus Soft Imaging System GmBH, Muenster). 20 photographs per WM tract (anterior, lateral and dorsal columns) and per animal were taken at 10,000X magnification. Blind analysis of the G-ratio was performed using ImageJ as described ^70^.

### Electrophysiology

Propagated compound action potentials (CAPs) were measured as reported ^68^ in the fornix of 14-month-old wt (n=4) and Taf1d38 (n=4) mice. Axial 400 μm-thick sections were incubated in artificial cerebrospinal fluid (aCFS) (124 mM NaCl, 2.5 mM KCl, 10 mM glucose, 25 mM NaHCO_3_, 1.25 mM NaH_2_PO_4_, 2.5 mM CaCl_2_ and 1.3 mM MgCl_2_) for 1 h. CAPs were recorded with a pulled borosilicate glass pipette (≈ 1 MΩ resistance) filled with aCFS by stimulation of the fornix with a bipolar electrode (CE2C55 FHC) placed 1 mm from the recording electrode with up to 2 mA, 100 µs pulses (Master-8, AMPI). Latencies between the onset of the stimulus artifact and the two negative waves of the CAP’s (N1 and N2) were measured with pClamp 10.0 (Molecular Devices). At least 20 independent recordings for N1 and 10 for N2 per genotype were performed.

### Behavioural testing

*Open field test*. Locomotor activity was measured as previously described ^62^ in clear Plexiglas® boxes of 27.5cm x 27.5cm, outfitted with photo-beam detectors (MED Associates’ Activity Monitor). Mice were placed in the center of the box and left to move freely for 15 min. Total traveled ambulatory distance and number of vertical episodes when the mouse stands on its hindlimbs were measured.

*Limb clasping.* Assessment of corticospinal function was performed by the limb clasping test ^69^. Mice were held vertically by the tail for 10 s and recorded. Videos were blindly scored as follows: 0 – normal extension of both hindlimbs; 1 – unilateral hindlimb partial retraction during more than 50% of the recorded time; 2 – bilateral hindlimb partial retraction during more than 50% of the recorded time; 3 - bilateral hindlimb total retraction during more than 50% of the recorded time.

*Rotarod.* Motor coordination was assessed in an accelerating rotarod apparatus (Ugo Basile). Mice were trained in 2 trials of 1 min at 4 rpm and 2 trials of 2 min with acceleration from 4 to 8 rpm over 1 min and the second minute at 8 rpm. The test was performed the following day with the rotarod set to accelerate from 4 to 40 rpm over 5 min in 4 independent trials. The latency to fall from the rod was measured as the mean of the four trials.

### Primary mixed culture and treatment of mouse cortical tissue

Cultures of mouse cortical neurons + glia from E17 wt brain cortex were prepared as described ^70^. Briefly, cerebral cortices were mechanically dissociated in MEM (ThermoFisher Scientific Gibco), supplemented with 10% horse serum, 22.2 mM glucose, 2 mM Glutamax-I (ThermoFisher Scientific Gibco) and penicillin-streptomycin (Sigma). The tissue was incubated in a 0.25% trypsin and 1 mg/mL DNase (Roche and dissociated using fire polished Pasteur pipettes. Cells were plated at a density of 80,000 cells/cm2 on plates or coverslips previously treated with poly-l-lysine (100 μg/mL) and laminin (4 μg/mL). Medium was replaced after 2 h by Neurobasal medium containing B27 supplement, 2 mM Glutamax-I (ThermoFisher Scientific Gibco) and penicillin-streptomycin. Glutamax-I was removed at DIV4, and at DIV7 cells were treated with either vehicle-1 (DMSO) or 2 µM CA074Me (Sigma, 205531). 1 h later, either vehicle-2 (100% ethanol) or 40 µM sphingomyelin (Santa Cruz, sc-201381) were added, and 8 h later the cells were harvested. Protein extracts were prepared as described in Western Blot section. For immunofluorescence, cells were fixed with 4% PFA and 4% sucrose and permeabilized with 0.1% Triton X-100 in PBS. After blocking in 2% BSA, 1% FBS and 0.1% Triton X-100 in PBS, the coverslips were incubated overnight at 4 ⁰C with anti-CTSB (1:400; RD systems, AF965) and rat anti-Lamp1 (1:400; DSHB, clone 1D4B), incubated with Alexa-conjugated secondary antibody and counterstained with DAPI. Images were acquired on a Zeiss LSM710 confocal vertical system. Mean gray value and integrated density were measured with Fiji and the Mander’s coefficient was determined using the JACoP plugin (Fiji).

### Plasmid generation

Human TAF1 cDNA (TAF1FL, ENST00000373790) was cloned into a pcDNA3 plasmid and EGFP sequence was added N-terminally by restriction free cloning ^71^ (RFC) to generate the pcDNA3-EGFP-TAF1FL plasmid. Exon 38 sequence was deleted and substituted by intron 37 retention sequence until the stop codon to generate the pcDNA3-EGFP-TAF1d38 plasmid. pEGFP-C1 was used as control plasmid. For TAF1 antibody validation, pcDNA3-TAF1FL without EGFP was cloned by RFC.

### Cell line culture and transfection

Oli-neu cells were grown in DMEM supplemented with 1% horse serum (Gibco), 10 μg/mL insulin (Sigma I9278), 10 μg/mL apotransferrin (Sigma T4382), 100 μM putrescine dihydrochloride (Sigma P5780), 200 nM progesterone (Sigma P0130), 220 nM sodium selenite (Sigma S5261) and penicillin-streptomycin (Sigma) at 37 °C with 10% CO2. The cells were transfected with either pEGFP-C1, pcDNA3-EGFP-TAF1FL or pcDNA3-EGFP-TAF1d38 using jetOPTIMUS transfection agent (Polyplus; 1.5 µL/µg of plasmid). Technical triplicates were carried for each condition, and cells were harvested after 24h.

293T cells were grown in DMEM supplemented with 10% FBS (Gibco) and penicillin-streptomycin (Sigma) at 37 °C with 5% CO2. The cells were transfected with either pcDNA3 or pcDNA3-TAF1FL using LipofectamineTM 2000 (ThermoFisher; 0.75 µL/µg of plasmid), or TAF1 siRNA against exon 32 sequence (GCCAACAGUGUUAAGUACAAU, Sigma) or scramble siRNA (Dharmacon Reagents) using RNAiMAX (ThermoFisher; 0.8 µL/µL of siRNA), and were harvested after 48h.

### GFP-Trap

Oli-neu cells were scrapped in PBS with protease inhibitors (PBS+PI: 1mM PMSF, Phosphatase Inhibitor Cocktail 3 [Sigma-Aldrich, P004] and Complete [Roche]) and centrifuged for 3 min at 500g at 4°C. After a washing in PBS+PI, the samples were incubated in lysis buffer (50 mM NaCl, 20 mM TrisHCl pH 8, 0.5% Triton X-100, 2 mM MgCl_2_) with 0.1 U/μL of benzonase for 30 min in ice. 250 mM NaCl, 0.5 mM EDTA and 0.1 mM DTT were added and after 5 min centrifugation at 16,000g at 4°C, the supernatant was collected, saving 10% of the volume as input. GFP-Trap dynabeads (Chromotek) were washed with IP buffer (150 mM NaCl, 0.5 mM EDTA, 20 mM TrisHCl pH 7.5) supplemented with protease inhibitors, and each sample was incubated with 25 μL of beads in IP buffer for 1 h at 4 °C. After 3 washes with IP buffer, beads were stored at-20 °C until usage.

### Mass spectrometry and proteomic data processing

Identical volumes of each immunoprecipitate were loaded on STRAP columns (PROTIFI, Farmingdale) and digested with trypsin in reducing and alkylating conditions. Tryptic peptides were desalted using C18 tips and subjected to independent LC-MS/MS analysis using a nano liquid chromatography system (Ultimate 3000 nano HPLC system, Thermo Fisher Scientific) coupled to an Orbitrap Exploris 240 mass spectrometer (Thermo Fisher Scientific). 5 µL of each sample were injected on a C18 PepMap trap column (5 µm, 300 µm I.D. x 2 cm, Thermo Scientific) at 20 µL/min, in 0.1% formic acid (FA), and the trap column was switched on-line to a C18 PepMap Easy-spray analytical column (2 µm, 100 Å, 75 µm I.D. x 50 cm, Thermo Scientific). Equilibration was done in mobile phase A (0.1% FA), and peptide elution was achieved in a 90 min gradient from 4%-50% B (0.1% FA in 80% acetonitrile) at 250 nL/min. Data acquisition was performed in data dependent acquisition, full scan positive mode. Survey MS1 scans were acquired at a resolution of 60,000 at m/z 200, scan range was 375-1200 m/z, with Normalized Automatic Gain Control (AGC) target of 300% and a maximum injection time of 45 ms. The top 20 most intense ions from each MS1 scan were selected and fragmented by higher-energy collisional dissociation of 30%. Resolution was set to 15,000 at m/z 200, with AGC target of 100 % and maximum ion injection time of 80 ms. Precursor ions with single, unassigned, or more than six charge states from fragmentation selection were excluded.

Raw files were processed using Proteome Discoverer v2.5 (Thermo Fisher Scientific). MS2 spectra were searched using Mascot v2.7 against a target/decoy database built from sequences corresponding to the *Mus musculus* reference proteome from Uniprot Knowledge database including contaminants. Search parameters considered fixed carbamidomethyl modification of cysteine, methionine oxidation, possible pyroglutamic acid from glutamine at the peptide N-terminus and N-terminus acetylation. The peptide precursor mass tolerance was set to 10 ppm and MS/MS tolerance was 0.02 Da, allowing for up to two missed tryptic cleavage sites. Spectra recovered after FDR ≤ 0.01 filter were selected for quantitative analysis. Label-free protein quantification used unique + razor peptides. Proteins with an adjusted p-value<0.05 and a log2(fold-change)>|3| respect to EGFP were defined as interactors. Proteins with an adjusted p-value<0.05 and a log2(fold-change)>|0.6| in the comparison between TAF1FL and vs TAF1d38 were defined as differential interactors. MAGMA gene-based analysis was performed for genes encoding the TAF1FL>TAF1d38 interactors. Note that GTF2H4 is located on the complement region on Chr6, so its genetic association might be not as robust as those found for GTF2B and FLII.

### Total RNA isolation and cDNA synthesis

RNA was isolated from striatum of 2-month-old wild type (n=4) and Taf1d38 (n=4) mice and from NAG+WM cortex of control subjects (n = 10) and individuals with PPMS (n = 10) or SPMS (n=10). Mice were subjected to daily manipulation for one week prior to sacrifice in their environment, to avoid any stress that could modify the brain basal transcriptomic state. Total RNA was extracted using the Maxwell 16 LEV simplyRNA Tissue Kit (Promega, AS1280). RNA quantification and quality were performed on a NanoDrop One spectrophotometer (Thermo Fisher) and an Agilent 2100 Bioanalyzer. For mouse samples, an RNA integrity number (RIN) over 8.5 was required, while for human samples the RIN cut-off was set to 6 for RNA-seq, and to 5 for RT-PCR analyses.

500 ng of total RNA was retrotranscribed using the iScript cDNA Synthesis kit (Bio-Rad, PN170-8891) with the following thermal conditions: 5 min at 25 °C; 20 min at 46 °C and 1 min at 95 °C.

### Real-time q-PCR

Quantification of more abundant events (total TAF1 (exons 25-28), intron 37 retention and complete exon 38) was performed using a CFX 384 Real Time System C1000 Thermal Cycler (Bio-Rad) in combination with SsoFast Eva Green (Bio-Rad, CN 172-5204), using 0.5 μM of each primer. Primers (table S20) were tested against isoform-specific amplicons by q-PCR, and PCR assay conditions were adjusted to obtain a single peak in the melting curve. ValidPrime kit (Tataa Biocenter PN A105S10) was used as control for genomic background, and technical triplicates were performed. Data were analyzed by GenEx 7.1.1.118 software (Multid AnaLyses AB). Relative quantification was performed (2-Δ(ΔCq) method) and mRNA levels were normalized relative to ACTB, GAPDH and TUBB in each sample.

### Digital-droplet PCR

Quantification of less abundant events (cryptic intron in exon 38 and distal exon 1) as well as total Taf1 (exons 25-28) that was used for normalization, was performed using a QX200 AutoDG Digital Droplet PCR system (Bio-Rad). Primers (table S20) were tested against isoform-specific amplicons by dd-PCR, and optimal annealing temperatures for each primer pair were determined through a temperature gradient from 55 °C to 65 °C.

PCR reactions were performed in triplicates with ddPCR Eva Green Supermix (Bio-Rad, CN 1864034), 0.25 μM of each primer and between 2.5 and 40 ng of template cDNA, depending on the relative abundance of the events. The reaction mix was combined with Automated Droplet Generation Oil for EvaGreen (Bio-Rad CN. 1864006) to generate the droplet emulsion using the Bio-Rad QX200 droplet generator, and processed in C1000 Touch Thermal Cycler (Bio-Rad). Droplets were read on the QX200 Droplet Reader (Bio-Rad, 1864003) and data were analyzed using QuantaSoft software v1.7.4.0917 (Bio-Rad). The absolute quantification of target DNA copies was performed by measuring the number of positive and negative droplets, and the copies/ng RNA were calculated. Finally, normalization relative to total Taf1 copies was performed.

### Bulk RNA sequencing

For mouse samples (n = 4), libraries were polyA+, had a read length of 1 x 50 bp, and were sequenced in HiSeq2500, generating >45 million single-end reads per sample. For human samples, libraries were polyA+, with a read length of 2 x 150 bp and were sequenced in NovaSeq6000, generating >210 million paired-end reads per sample.

### Bulk RNA-seq data processing and analysis

Quality analyses were performed using FastQC 0.11.9 software. Salmon 1.9.0 alignment mapping algorithm was used ^72^, employing indexes constructed with either *Mus musculus* reference genome (Ensembl GRCm39) or *Homo sapiens* reference genome (Ensembl GRCh38). Differential expression analysis at gene level was carried with DESeq2 1.40.2 ^73^. Genes with less than 20 normalized counts on average in both wt and Taf1d38 groups or CT and PPMS groups were filtered out. Significant differentially expressed genes (DEGs) were defined for an adjusted p value < 0.05.

DEGs were validated *in silico* via gene-based association test via MAGMA 1.9 ^34^, using the GWAS summary statistics from the International Multiple Sclerosis Genetics Consortium MS (47,429 MS cases; 68,374 controls) ^6,7^. For MAGMA analysis the model applied was ‘multi=snp-wise’, which aggregates the results of the ‘snp-wise=top’ and ‘snp-wise=mean’ analyses models, improving the power and accuracy of gene-based association tests especially when the genetic model of the disease under study is unknown. For gene-based analysis, gene boundaries were retrieved from RefSeq (GRCh38.p14), lifted over to GRCh37 coordinates, and expanded to 5 kb flanking regions, to encompass potential regulatory elements.

Principal component analysis with medium to highly expressed genes (> 150 counts) in human samples revealed that one of the controls (CT4) grouped together with the 3 PPMS samples (fig. S6A), thus, we decided to remove CT4 from the DEG analysis. Clustering of the remaining CT vs PPMS samples resulted as expected (fig. S6B) and differential expression analysis was performed in this subset.

TAF1 alternative splicing was explored using three complementary software: 1) vast-tools v. 2.5.1 ^29^ using human junction libraries for the align and combine modules, and a |dPSI| (absolute difference in per cent spliced in) ≥ 10 in the compare module; 2) rMATS v.4.1.2 ^74^ using STAR v. 2.7.10 ^75^ to align fastq reads together with a custom splice-junction library, and a FDR < 5% for differential splicing analysis; and 3) Majiq v. 2.5 ^76^ after alignment with STAR and default parameters, using both deltapsi and heterogen, with a probability of changing >95% The STAR aligned reads were also processed with samtools 1.9 and the profiles were visualized using the Integrative Genomics Viewer (IGV) 2.4. Sashimi plots at the 3’ end of TAF1 gene were also generated with IGV 2.4.

### Overlap analyses

http://nemates.org/MA/progs/overlap_stats.html was used. It runs an hypergeometric test and provides a representation factor, which is the number of overlapping genes divided by the expected number of overlapping genes drawn from two independent groups. Common DEG signatures were obtained by comparing polyA+ Taf1d38 mice and PPMS NAG+WM datasets.

### Single nuclei RNA sequencing

2-month-old wt (n = 3) and Taf1d38 (n = 3) mice were manipulated as described for bulk RNA-seq experiment, while 10-month-old wt (n = 3) and Taf1d38 (n = 3) mice were sacrificed without previous manipulation.

The 2-month-old samples were processed in two batches (first batch n = 2, second batch n = 1) using the whole striatum, while the 10-month-old samples were processed all together using the right half of the spinal cord. The samples were homogenized in nuclei isolation buffer (10 mM Tris-HCl pH 8.0, 0.25 M sucrose, 5 mM MgCl_2_, 25 mM KCl, 0.1% Triton X-100, 0.1 mM DTT) and 0.2 U/μl RNase inhibitors (RNAsin Plus), sequentially filtered through a 70-μm and a 30-μm strainer and centrifuged for 10 min at 900g at 4 °C. The pellet was resuspended in 150 μl of nuclei isolation buffer and mixed with 250 μl of 50% iodixanol for debris removal. 500 μl of 29% iodixanol were overlaid with this solution, and centrifuged for 30 min at 13,500g at 4 °C. The pellet was resuspended in 100 μl PBS with 1% BSA and 0.2 U/μl RNase inhibitors.

The nuclei were stained with Trypan Blue 0.4% and counted in an haemocytometer. A total of 10,000 estimated nuclei for each sample were loaded on the Chromium Next GEM Chip G, although a much lower number of nuclei was recovered after sequencing in the 2-month-old experiment. cDNA libraries were prepared following the Chromium Next GEM Single Cell 3′ Reagents Kits v3.1 User Guide and sequenced in NovaSeq6000 (2-month-old samples) or NovaSeq X Plus (10-month-old samples).

### Single nuclei RNA-seq data processing and analysis

The samples were aligned with Cellranger v.7.0.1 with reference genome GRCm38-mm10. Filtered count matrices and barcodes were uploaded into R 4.3.2 to be processed following Seurat v4 and v5 ^77^. The 2-month-old samples were processed separately for initial QC to avoid differences related to sequencing batches. Cells were filtered for number UMIs per nucleus, number of genes per nucleus and mitochondrial percentage (2-month-old samples: percent_mito < 2 & nFeature_RNA > 50 & nFeature_RNA < 5000 & nCount_RNA > 5 & nCount_RNA < 10000; 10-month-old samples: percent_mito < 1 & nFeature_RNA > 250 & nFeature_RNA < 6000 & nCount_RNA > 200 & nCount_RNA < 12000). The final selected nuclei had the following median number of UMI, median number of genes and median mitochondrial percentage: 2-month-old samples (wt: 8056, 3376, 0.1804958; Taf1d38: 8404, 3454, 0.1160056) with 10,351 final nuclei; 10-month-old samples (wt: 644, 523, 0.1; Taf1d38: 566, 461, 0.1) with 47,649 final nuclei.

Log-normalization, feature selection and SCT normalization regressing mitochondrial percentage, were performed in the whole object with the 6 samples in each experiment. The first 30 PCs were selected using variable features determined by *SCTransform* (2000 genes). Sample integration was performed with Harmony ^78^ (*RunHarmony*) on the selected PCs using grouping variables the replicates and batch information. Dimension reduction was performed using *runUMAP*. KNN graph was constructed using *Findneighbors* with annoy method, 50 trees and K=20. Clusters were identified with *Findclusters* to different resolutions from 0.2 to 1.2.

In the 2-month-old dataset, first exploratory view of the clusters showed expression hormonal genes (“Prl”, “Gh”, “Pou1f1”, “Tshb”) from nuclei corresponding only to one sample, which are not expected to be expressed in the striatum, and probably came from contamination from the hypothalamus or the bed nucleus of the stria terminalis. An “Hormonal_gene_score” was calculated using *AddModuleScore* with 4 controls and 50 bins for all the nuclei, and we filtered out the possibly contaminated nuclei by “Hormonal_gene_score”>0. All the processing from the QC to cluster identification was repeated without these spurious cells.

Clusters were annotated using *FindAllMarkers* based on classical top markers per cluster. In the 2-month-old experiment, annotation was complemented with label transfer using the dataset from Muñoz-Manchado *et al*. ^36^. The GSE97478 annotated expression matrix was downloaded from GEO repository, and common anchors were calculated (*FindTransferAnchors*, with parameters cca reduction and SCT normalized query and reference). Annotations were transferred with *TransferData* on the first 30 dimensions of the cca reduction to the query dataset. In the spinal cord samples, oligodendrocyte populations annotation was complemented with label transfer using the dataset from Marques *et al*. ^33^.

Differential expression analysis was performed in each annotated cell type and running *FindAllMarkers* between wt and Taf1d38 cells using Wilcoxon Rank Sum test. As a complementary analysis, differential gene expression was run in pseudobulk in the annotated clusters using EdgR ^79^.

### Identification of perturbed cell types

AUGUR R package ^80^ (https://github.com/neurorestore/Augur) was used to identify perturbed cell types through a machine-learning model that predict the condition (wt or Taf1d38) of each cell type and reports an AUC score of the accuracy of the model. AUC score > 0.5 was considered as a significant perturbation after running different iterations by downsampling to the same number of nuclei per comparison.

### Pathways and Gene ontology categories

Each nuclei from spinal cord oligodendroglia clusters was scored using GSVA Bioconductor package ^81^. The average score per annotated cell type (Hallmarks and GO category) was calculated with AverageExpression, ranging from 0 to 1. Heatmaps were built with ComplexHeatmap R package ^82^.

### Functional enrichment analysis

The Ingenuity Pathway Analysis software (QIAGEN Inc., v.107193442) was used to determine enriched canonical and disease pathways within Taf1d38 DEG signatures, TAF1FL>TAF1d38 interactors (after eliminating potential unspecific interactors such as cytoskeletal proteins) and genes with top altered RNAPII dynamics in Taf1d38 mice.

DEGs were also examined for enriched categories via GSEA with clusterProfiler v.4.8.1 package in R ^83^, using a modified Kolmogorov-Smirnov statistics to assess significant categories. Enriched categories were validated for MS with a gene-set level analysis via MAGMA using Z scores from gene p-values.

### Bulk ChIP sequencing

Mice were manipulated as described for bulk RNA-seq experiment. A pool of 4 striata was used for each sample, and a total of 2 wt and 2 Taf1d38 pooled samples were processed in 2 batches. Chromatin immunoprecipitation was performed as previously described ^84^. After homogenizing the tissue in ice-cold PBS+PI, the extracts were crosslinked with 1% formaldehyde for 10 min at 37 °C, and the reaction was quenched with 125mM glycine. After 5 min centrifugation at 5,000g at 4 °C and washing in ice-cold PBS+PI, the pellets were resuspended in lysis buffer A (5 mM Pipes pH 8.0, 85 mM KCl, 0.5% NP40) supplemented with protease inhibitors and incubated for 10 min on ice. Cell nuclei were pelleted by 5 min centrifugation at 4,000g at 4°C, and resuspended in lysis buffer B (50 mM Tris HCl pH 8.1, 1% SDS, 10 mM EDTA) supplemented with protease inhibitors. Chromatin was sonicated for 25 min using a Covaris ultrasonicator (E220 Evolution) with the following parameters: 200 cycles/burst, 5% duty factor and 140 W of peak incident power. After a 10 min centrifugation at 20,000g the supernatant was collected. 50 μL of sonicated chromatin was reverse-crosslinked using Proteinase K in lysis buffer B at 65°C overnight and, after phenol chloroform extraction, DNA fragmentation was checked (intended fragment size 200-600 bp). For each immunoprecipitation (IP), 40 μg of chromatin spiked-in with 5% of human chromatin from MCF7 cell line, and 4 μg of antibody (rabbit anti-Rpb1, Cell Signaling, Cat.14958; rabbit anti-RNA Pol II phospho S2, Abcam, ab5095; rat anti-RNA Pol II phospho Ser5, Sigma, MABE954) was incubated in IP buffer (0.1% SDS, 1% Triton X-100, 2mM EDTA, 20 mM TrisHCl pH8, 150 mM NaCl) at 4 °C overnight. 10% of chromatin in IP buffer was stored as input. Dynabeads protein A and G (ThermoFisher) were blocked overnight with IP buffer with 1 mg/ml BSA, and each IP was incubated with 25 μL of beads for 4 hours at 4 °C. Beads were sequentially washed with buffer 1 (20 mM Tris HCl pH 8, 0.1% SDS, 2 mM EDTA, 150 mM NaCl), buffer 2 (20 mM Tris HCl pH 8, 0.1% SDS, 2 mM EDTA, 500 mM NaCl), buffer 3 (20 mM Tris HCl pH 8, 1% NP40, 1% sodium deoxycholate, 1 mM EDTA, 250 mM LiCl) and T buffer (10 mM Tris HCl pH8), all supplemented with protease inhibitors. ChIPmentation on beads was performed as previously described ^85^, using Tn5A enzyme provided by the Proteomics Service of Centro Andaluz de Biología del Desarrollo in tagmentation buffer (10 mM Tris HCl pH8, 10% dimethylformamide, 5 mM MgCl_2_) for 10 min at 37 °C. After serial washes in buffer 3 and TE buffer (10 mM Tris HCl pH8, 1mM EDTA), tagmented DNA was eluted with 1% SDS and 100 mM NaHCO_3_ in TE buffer at 50 °C for 30 min. Overnight decrosslinking was performed using Proteinase K in 175 mM NaCl at 37 °C. After incubation with RNase A (ThermoFisher), DNA was purified (QIAGEN, 28106) and inputs were tagmented and purified. Libraries were amplified for the optimum cycles determined by qPCR using NEBNext High-Fidelity Polymerase (New England Biolabs, M0541), purified with Sera-Mag Select Beads (GE Healthcare, 29343052) and sequenced after Qubit quantification and size profile analysis with Agilent 2100 Bioanalyzer, using Illumina NextSeq 500 and single-end configuration with a read length of 75 bp.

### ChIP-seq data processing and analysis

FASTQ Toolkit 1.0.0 and Trimmomatic 0.36 (20:4 sliding window, 2:leading, 2:trailing, 35:minlen) were run for trimming. The reads from the IP and input libraries were aligned to the mouse and human reference genomes using bowtie 2.4.5, and signal tracks were obtained with bamCoverage (deepTools 3.5.1 ^86^).

To correct technical biases and global differences, spike-in normalization was applied using human chromatin as control. RPGC-normalised tracks were subsequently scaled by the occupancy ratio (OR), which was calculated as follows: from raw signal tracks, enrichment (IP/input) was calculated and averaged for all the positive (i.e. counts in both IP and input sample) 50 bp bins. OR was then obtained by dividing average enrichment in human and mouse reads as OR = average(Eh)/average(Em), where Eh = enrichment at each bin of the human experimental genome and Em = enrichment in each bin of the reference mouse genome. In each case, the enrichment is calculated as E= C^IP^ / C^IN^ where C^IP^ = counts in the IP sample at each genome bin and C^IN^ = counts in the input sample at each genome bin. BedCoverage (Samtools v1.6.) was used for quantification at regions of interest using BAM files. CPM values were scaled by OR to allow inter-sample comparisons. Spike-in quantifications were used to generate normalized RPGC counts tables and signal profiles with Seqplots ^87^.

For signal distribution analysis, two regions were defined: the promoter region, which comprised the sequence 1.5 kb up and downstream of the transcription start site (TSS); and the gene body region, defined between the TSS and the transcriptional termination site (TTS). Only genes larger than 4.5 kb were plotted in total RNAPII and Ser5P RNAP II metaplots, as shorter genes may introduce noise in the promoter/gene body definition. Genes larger than 30 kb were plotted in Ser2P RNAPII metaplots, and the quantification shown in box plot was performed between 300 bp downstream the TSS and the TTS. Log2FC of Taf1d38 vs wt counts for each IP and each experiment were calculated. The top 15% genes with higher log2FC for total RNAPII or Ser5P RNAPII, or lower log2FC for Ser2P RNAPII in a given experiment were extracted. The overlapping genes that resulted from crossing the 15% top lists obtained in batches 1 and 2, were taken as the top affected genes. Thus, we obtained 1,285 genes with highest total RNAPII promoter region occupancy, 990 genes with highest Ser5P RNAPII promoter region occupancy and 2,397 genes with lowest Ser2P RNAPII gene body occupancy (table S18) in Taf1d38 mice. For functional enrichment analyses, a fold-change cut-off was defined in each case: FC>500 for RNAPII promoter occupancy (317 genes); FC>16 for Ser5P RNAPII promoter occupancy and log2(pausing index)> 2.5 (120 genes); and FC<0.7 for Ser2P RNAPII gene body occupancy (624 genes).

To estimate the level of paused RNAPII, we calculated the pausing index ^84^ as the ratio between the occupation of RNAPII within the promoter (computed as the sum of reads in 400 bp surrounding the TSS), and the occupation of RNAPII within the gene (calculated as the average number of reads in 400 bp windows throughout the gene body, starting 200 bp downstream the TSS). To ensure comparability with promotor occupancy analyses, only genes larger than 4.5 kb were considered (table S18).

Finally, metagene and pausing index measurements were performed with different subsets of genes according to mean total RNAPII (fig. S7C) or Ser2P RNAPII (fig. S7D) gene body occupancy quartiles in wt samples. The global trends observed with all genes in Fig. 6D were reproduced especially on the gene sets with higher RPGC counts.

### Statistical analysis

Statistics were performed with SPSS 21.0 (SPSS Statistic IBM). The normality of the data was checked by Shapiro-Wilk and Kolmogorov-Smirnov tests. For two-group comparison, two-tailed unpaired Student’s t-test or Mann-Whitney U test were performed. For multiple comparisons, one-way ANOVA test followed by a Tukey’s post hoc test or Kruskal-Wallis test were applied. A significance of p-value < 0.05 was used throughout the study. Benjamini-Hochberg correction was applied for multiple testing.

## Acknowledgements

Tissue specimens used in this research were obtained from the UK Multiple Sclerosis Tissue Bank at Imperial College (London) and the Netherlands Brain Bank. We thank the patients and families who participated in the tissue donation programs.

We thank Drs. Hauke Werner and Olaf Jahn (Max Planck Institute for Multidisciplinary Sciences, Göttingen, Germany) for advice on proteomics analyses of human MS tissue and Dr. Thomas Reinheckel (Institut für Molekulare Medizin und Zellforschung, Freiburg, Germany) for providing mouse model brain samples. We are grateful to Dr. M Dolores Ledesma and her team (CBM) for advice on lysosomal permeabilization protocols and oligodendroglial cell biology. We also thank Dr. Jorge R. Cabrera for advice on the role of TAF1 in brain diseases and Dr. Manuel Comabella (CIBER-NED) for advice on MS genetics, histopathology and therapeutics. We also thank Cristina Martos-Polo (Group of Neuro Immuno-Repair of the HNP) and Dr Javier Mazarío (Microscopy Service of the HNP) for their assistance with the histological quantifications in the human CNS and Nuria de la Torre (CBM) for administrative assistance.

Computational analyses were run at the Center for Scientific Computing provided by the Universidad Autónoma de Madrid (CCC-UAM), the High-Performance Computing cluster provided by the Centro Informático Científico de Andalucía (CICA) and the Swedish National Infrastructure for Computing (SNIC) at the Uppsala Multidisciplinary Center for Advanced Computational Science, partially funded by the Swedish Research Council through grant agreement no. 2018-05973.

We thank the following core facilities: CBM-CNB Mouse Transgenesis, CBM-Animal Facility, CBM-Genomics & Massive Sequencing, CBM-Bioinformatics, CBM-Advanced Light Microscopy Facility, CBM-Electron Microscopy, CNIO-Genomics Core Unit, CNIO-Proteomics Core Unit, Science for Life Laboratory-National Genomics Infrastructure in Stockholm and CRG-Genomics Unit in Barcelona.

CBM is a Severo Ochoa Center of Excellence (MICIN, Award CEX2021-001154-S).

## Author contributions

C.R.-L. contributed to study design and interpretation, and was involved in all assays and data collection.

I.H.H. contributed to study design and interpretation, and was involved in Taf1d38 mouse model generation and characterization. J.T-.B contributed to ChIP-seq and GFP-Trap experiments and analyses.

E.A. performed single nuclei RNA-seq analyses. J.P.-U. performed primary cultures and immunofluorescence experiments. D.L.-M. performed bioinformatics and Western blot analyses. I.R.-B., A.M.A., M.G.-B., B.S.-G. and M.S.-G. performed immunohistochemistry and Western blot. I.G.-O. and M.M.-J. performed GSEA and gene-based analyses from GWAS data. M.L.-S. performed behavioral tests on mice. M.C.O. and T.H.J.M. performed immunohistochemistry on human samples. I.M.D. performed cultures. J.C.C, Z.M. and F.P.-C. performed electron microscopy, immunofluorescence and fluoromyelin analyses. C.Z. and M.K contributed to single cell RNA-seq experiments. A.P.-S. performed electrophysiological recordings. M.C.-B. performed bioinformatics analyses of C-terminal TAF1 motifs and conservation. A.P. performed and interpreted proteomic analyses. J.P.-U., A.B., B.A., F.dC., N.L.F, J.J.M.H, C.T., C.M., D.C., F.C.-L. and G.C.-B. made intellectual contributions to experimental design and discussion. C.M., F.C.-L., G.C.-B and J.J.L. directed the study and designed experiments. J.J.L. wrote the paper with input from all authors.

## Data availability

RNA-seq and ChIP-seq data are deposited in the Gene Expression Omnibus (GEO) database under the accession numbers GSE255578 and GSE261661 (under restricted access until publication). The RNA-seq datasets already published that were reanalyzed here are available at the GEO database under the accession number GSE138614. Source data are provided with this paper.

Standard procedures and published codes were used for data analysis (see Methods).

## Supplementary figure legends

**Fig. S1.**
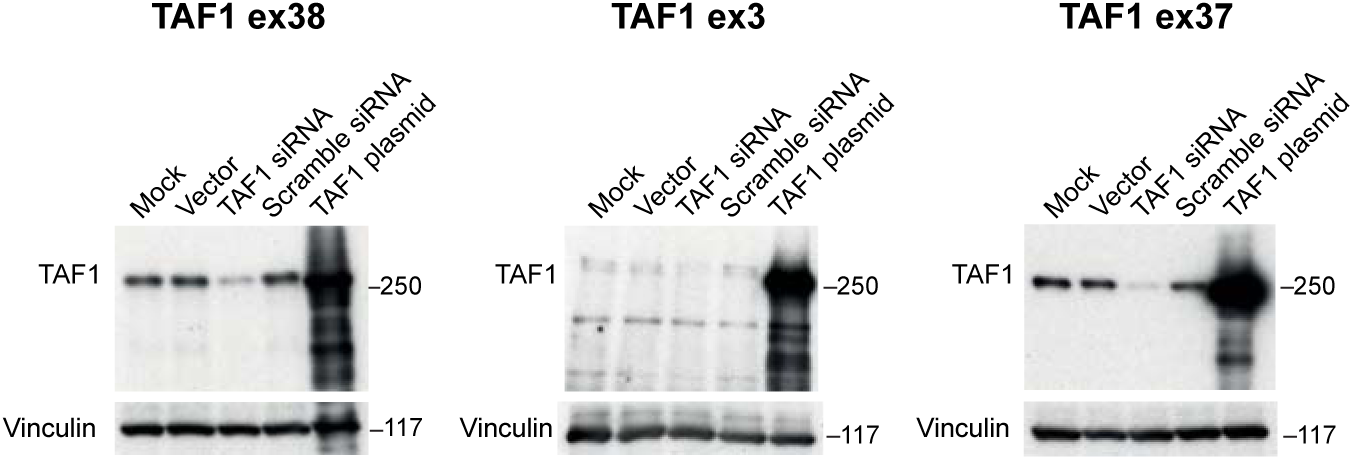
Validation of TAF1 antibodies and multiple sequence alignment of C-term TAF1. TAF1 detection with antibodies against sequences encoded by exons 38, 3 or 37 and vinculin levels in human 293T cells. Mock: transfection agent (lipofectamine); Vector: transfection with pcDNA3 plasmid; TAF1 siRNA: transfection with commercial siRNA against exon 32 sequence of human TAF1 mRNA; Scramble siRNA: transfection with a commercial scramble siRNA; TAF1 plasmid: transfection with pcDNA3-humanTAF1 plasmid.

**Fig. S2.**
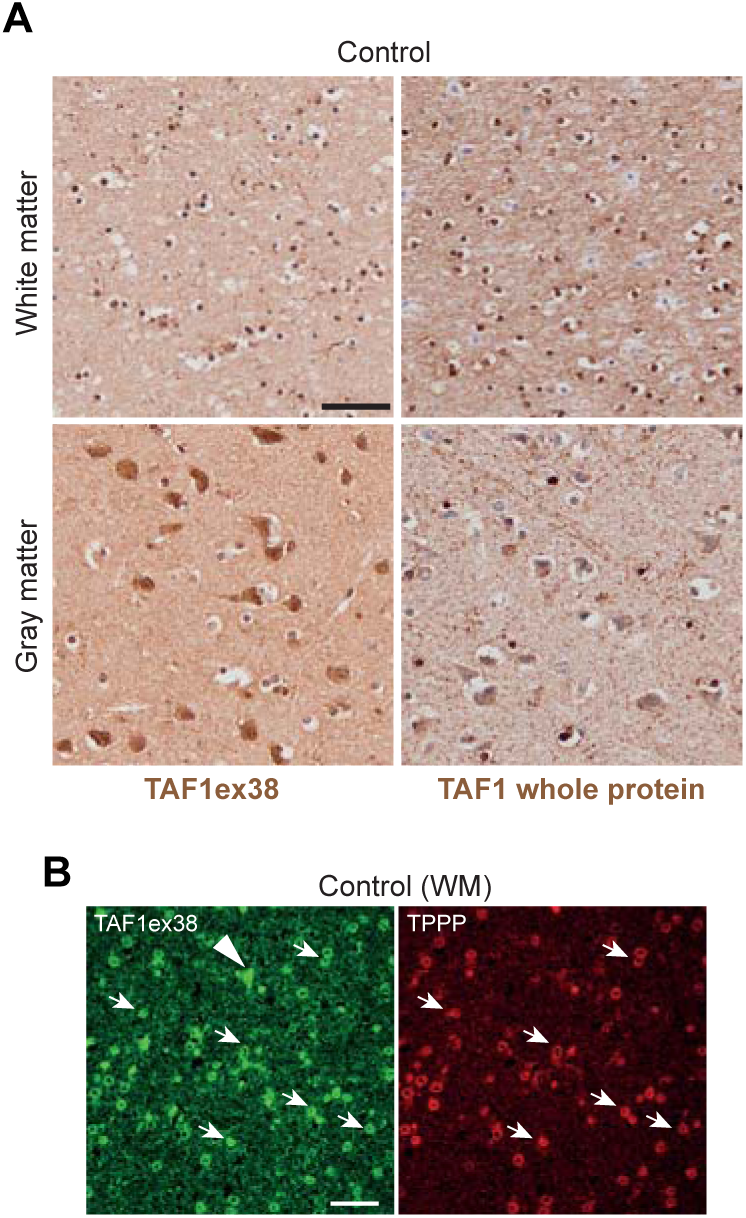
TAF1 staining pattern in control brain. (A) Immunohistochemistry in control cortical GM and WM with the TAF1ex38 antibody and the TAF1 antibody raised against the whole protein (TAF1 whole protein) with haematoxylin counterstaining. Scale bar: 50 µm. (B) Double immunofluorescence in control WM with TAF1ex38 antibody and TPPP, showing that most of the TAF1ex38+ cells express the oligodendrocyte marker TPPP (arrows), and few cells do not (arrowhead). Scale bar: 30 µm.

**Fig. S3.**
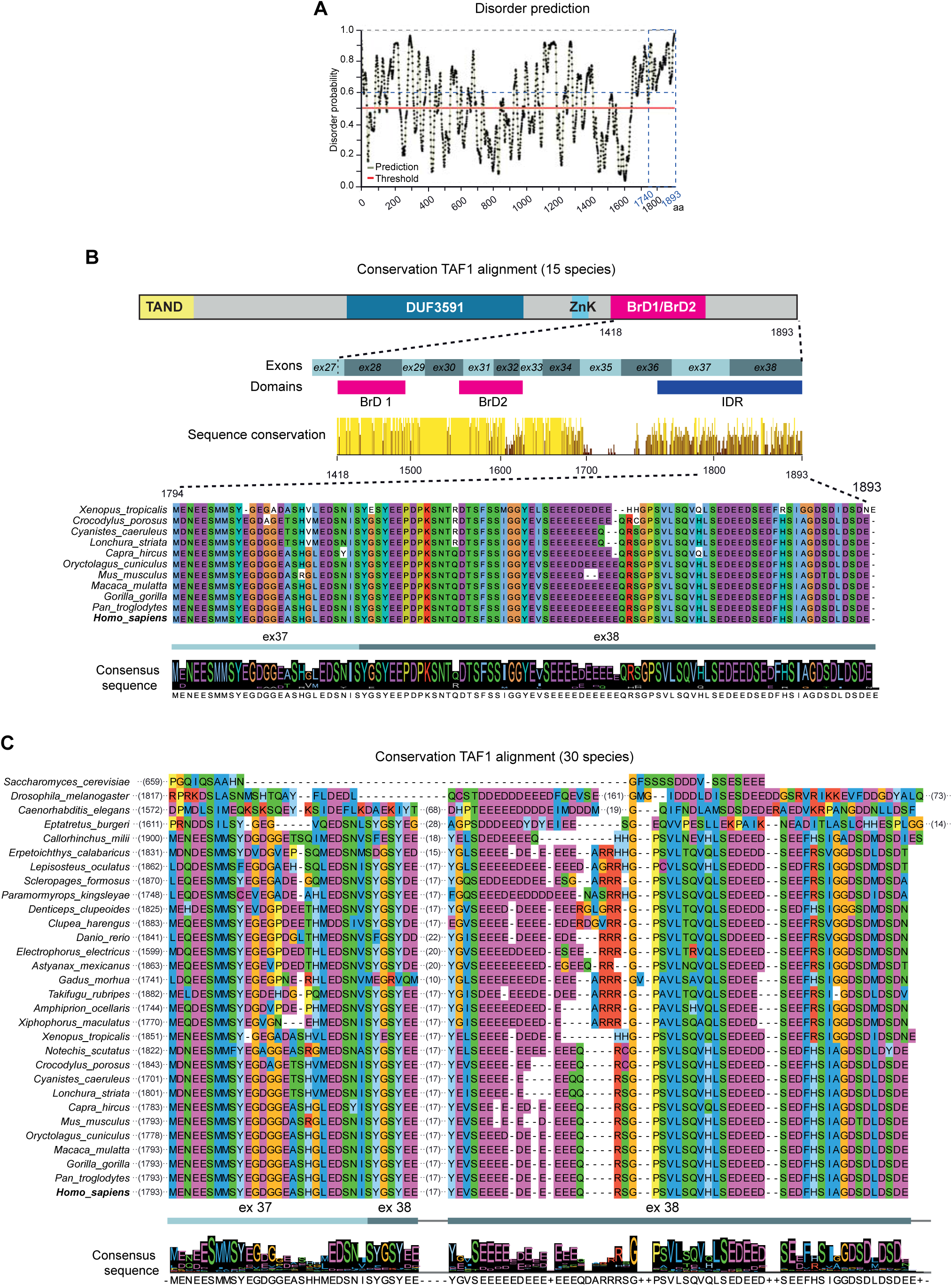
Multiple sequence alignment of C-term TAF1. (A) Disorder prediction of TAF1 determined with PrDOS webservice. Maximum false positive rate was established at 5% for the prediction, and a probability of 0.5 was taken as the threshold of disorder (red line). All amino acids in the C-term region (1740-1893 residues) have a disorder score higher than 0.6 (blue dashed line). (B) Conservation according to JalView of the last 476 amino acids of TAF1 protein (1418-1893 residues) across 15 species (the ones listed in the sequence alignment plus the reptile *Notechis scutatus* and the bony fishes *Takifugu rubripes*, *Xiphophorus maculatus* and *Amphiprion ocellaris*). The multiple sequence alignment corresponds to the 1794-1893 sequence encoded by part of exon 37 and the entire 38, residues are coloured according to Clustal X colour scheme. The protein consensus sequence is shown below. (C) Multiple sequence alignment also of the 1794-1893 sequence, but across 30 species (including *Saccharomyces cerevisiae, Drosophila melanogaster* and *Caenorhabditis elegans*) evidences that conservation extends beyond vertebrates. Note that interruptions in the alignment are caused by insertions in some sequences, which shift the rest in the alignment. The protein consensus sequence is shown below.

**Fig. S4.**
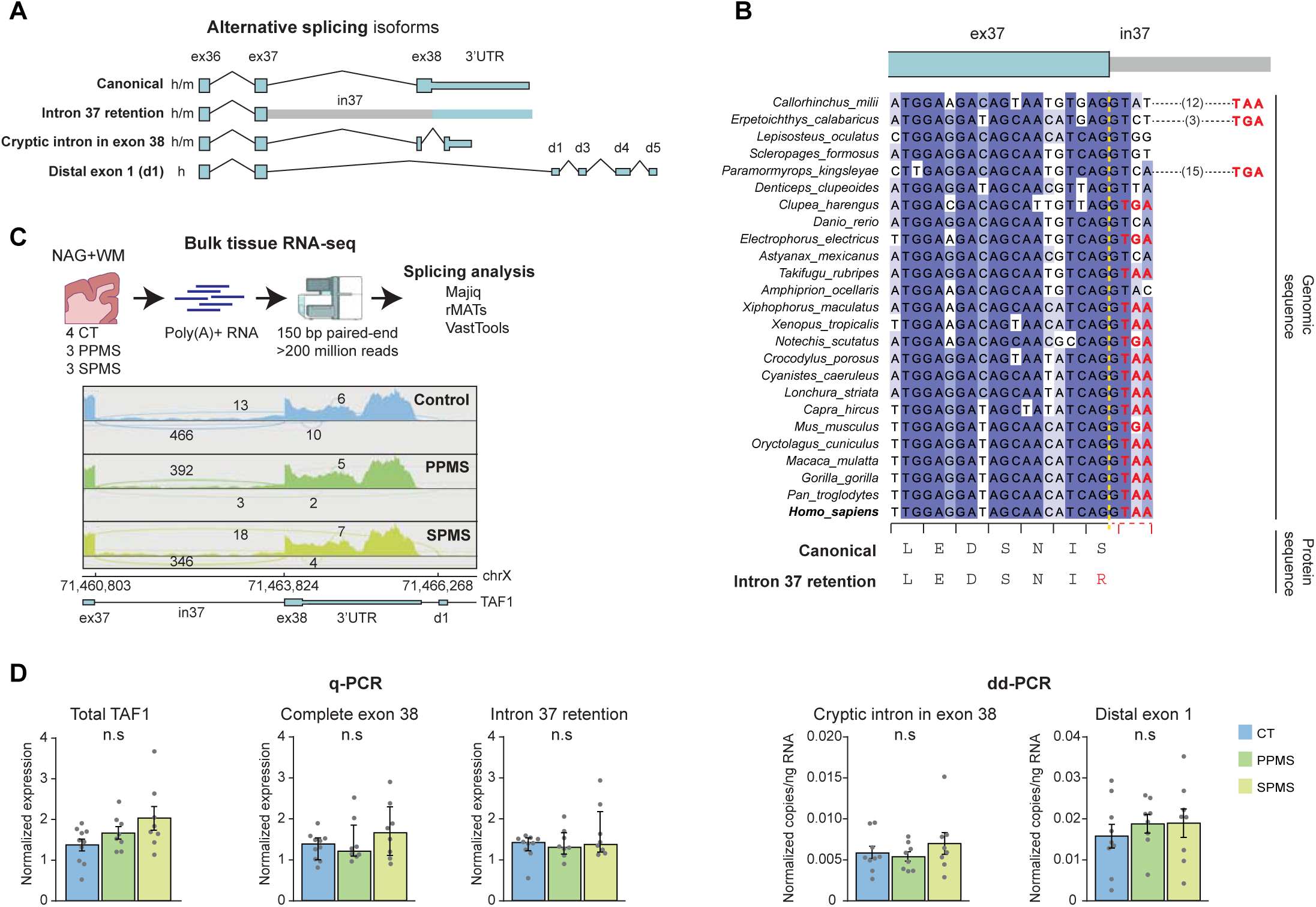
Analysis in MS brains of alternative splicing events affecting exon 38. (A) 3’-transcript structure of *TAF1* RNA isoforms. The *Canonica*l form includes exon 38 and the canonical 3’ UTR. The *Intron 37 retention* isoform (PSI=19 in cerebral cortex according to https://majiq.biociphers.org/majiqlopedia/) includes intron 37 and is predicted to generate a truncated protein lacking exon 38 due to a highly conserved stop codon (see B). The *Cryptic intron in exon 38* isoform (PSI<1 in cerebral cortex according to https://majiq.biociphers.org/majiqlopedia/) lacks the sequence encoding the last 52 amino acids and adds 21 C-terminal amino acids encoded by part of the canonical 3’ UTR. The *Distal exon 1* isoform (PSI=6 in cerebral cortex according to https://majiq.biociphers.org/majiqlopedia/) splices a large region comprising exon 38 and the 3’ UTR and incorporate distal exons. h: reported in human; h/m: reported in human and mouse. (B) Multiple sequence alignment of the genomic sequence comprising the last 20 nucleotides of exon 37 and the beginning of intron 37 across 25 species. A dashed yellow line separates the exonic from the intronic sequence. The amino acid sequence for *Homo sapiens* is shown below. Note the high conservation of the stop codon (red) if intron 37 is retained. (C) Bulk polyA+ RNA-seq on NAG+WM samples experimental design and workflow (up). Sashimi plots from IGV of the 3’ end of TAF1 transcripts in representative samples showing no differential splicing events affecting exon 38 inclusion. Drawings were imported from NIH Bioart. (D) Left: total TAF1, complete exon 38 and intron 37 retention mRNA levels normalized to housekeeping genes by q-PCR in NAG+WM from controls (n = 10), PPMS (n = 8) or SPMS (n = 8) individuals. Right: cryptic intron in exon 38 and distal exon 1 copies per ng RNA normalized to total TAF1 by digital droplet-PCR in NAG+WM from controls (n = 9), PPMS (n = 8) or SPMS (n = 8) individuals. Total TAF1 q-PCR and dd-PCR events: one-way ANOVA followed by Tukey’s post hoc test, graphs show mean ± SEM. Intron 37 retention and complete exon 38 inclusion events: Kruskal-Wallis test, graphs show median ± interquartile range. n.s., not-significant.

**Fig. S5.**
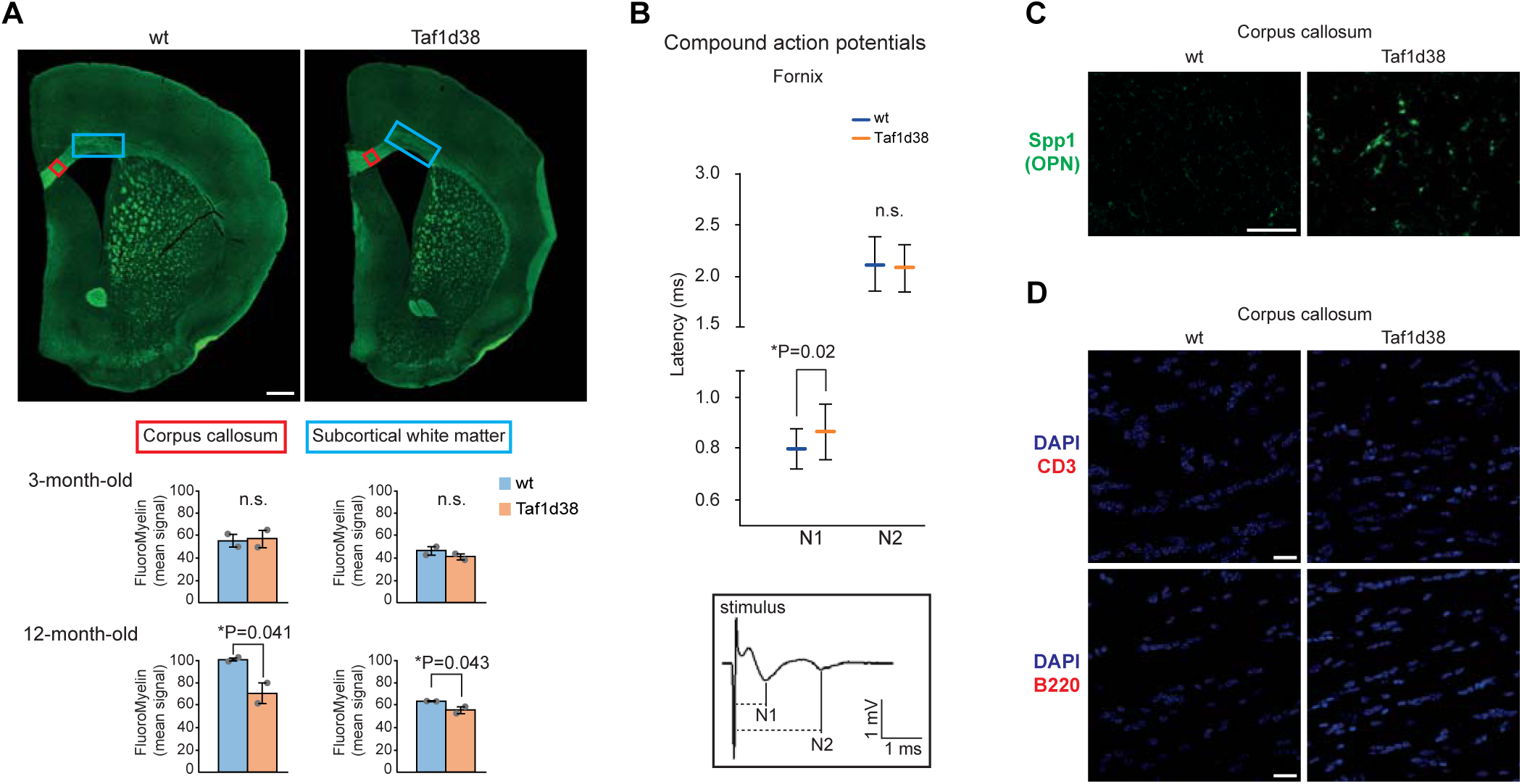
Additional histopathological characterization of Taf1d38 mice brain. (A) Examples of FluoroMyelin staining of coronal brain slices of 12-month-old mice (up). The boxes in corpus callosum and subcortical WM (cingulum bundle and medial external capsule) show the areas whose signal was quantified, as shown in the histograms (down). Quantifications of the mean intensity values were performed on 3 and 12-month-old wt and Taf1d38 mice (n = 2 for each age and genotype). One-tailed unpaired t-test. Scale bar: 500 μm. (B) Electrophysiological recording of propagated compound action potentials (CAPs) in the fornix of 14-month-old wt (n = 4) or Taf1d38 mice (n = 4), showing increased latency in the conduction through myelinated fibers. Latencies are defined as the time between the onset of the stimulus artifact and each of the two main negativities, N1 and N2, corresponding to conduction through myelinated and unmyelinated fibers respectively (see inset down). Two-tailed unpaired t-test. (C) Spp1 (osteopontin) immunofluorescence in corpus callosum (sagittal sections) from 6-month-old mice, exemplifying the upregulation in Taf1d38 WM. Scale bar: 50 µm. (D) CD3 or BC220 immunofluorescence together with DAPI nuclear counterstaining in corpus callosum (coronal sections) from 3-month-old mice, evidencing the absence of lymphocyte infiltration. Scale bar: 20 µm. (A,B) Graphs show mean± SD.

**Fig. S6.**
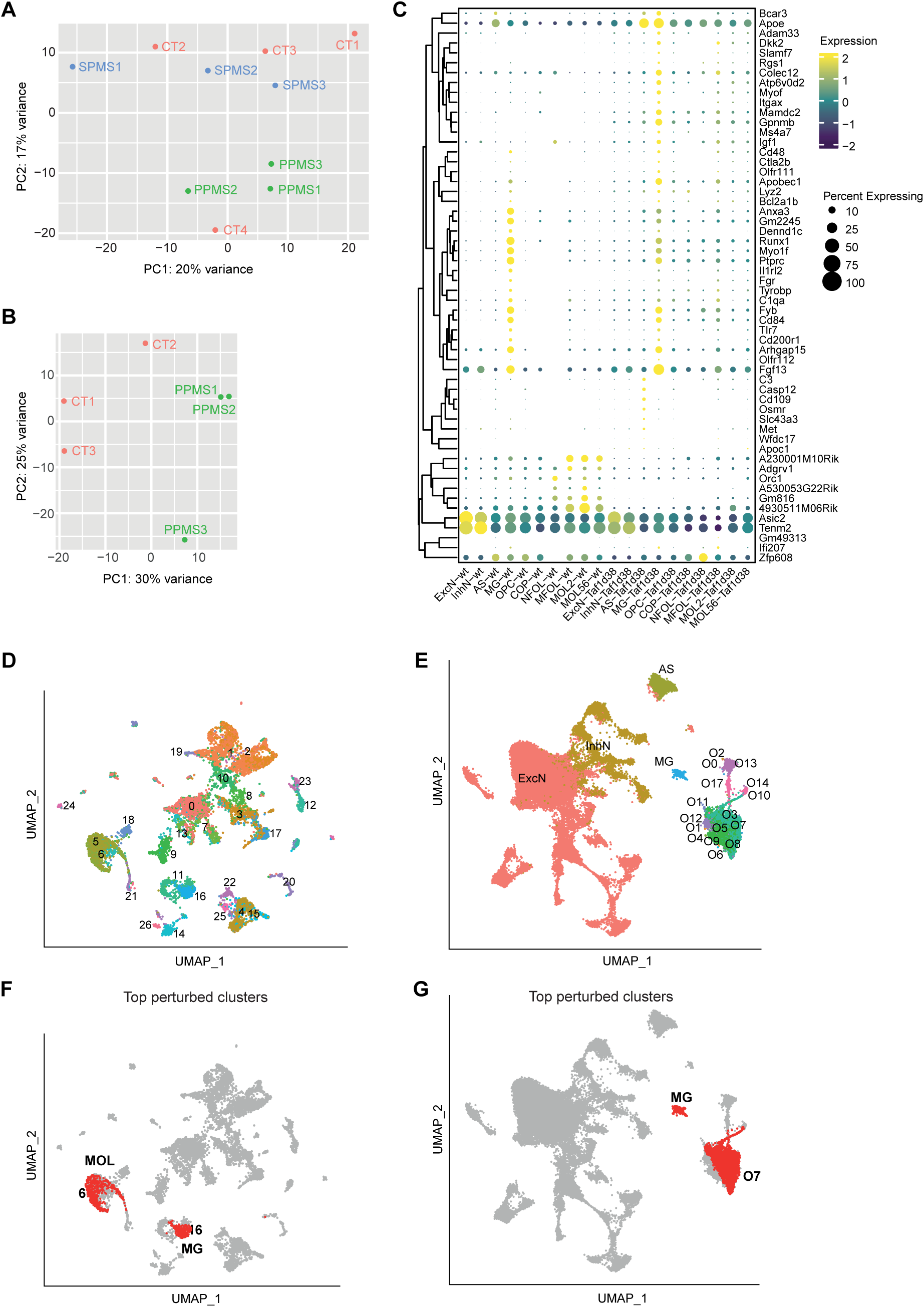
Additional information regarding bulk tissue RNA-seq on human samples and snRNA-seq on mouse samples. (A) Principal component analysis of genes in the polyA+ RNA-seq experiment on human cortical samples (n = 4 controls, n = 3 PPMS, n= 3 SPMS). The CT4 sample was atypical as it mapped closer to PPMS samples than to the rest of control samples. Accordingly, we decided to exclude this sample from the final analysis. (B) Principal component analysis of genes in the polyA+ RNA-seq experiment of the samples that were analyzed for DEG (n = 3 controls, n = 3 PPMS).(C), Dotplot of top DEG after pseudobulk analysis in clusters of spinal cord snRNA-seq. (D) UMAP plot of cell clusters in 2-month-old striatal snRNA-seq with resolution 1 yield a total of 27 clusters. (E) UMAP plot of cell clusters in 10-month-old spinal cord snRNA-seq with resolution 0.8 yield a total of 16 oligodendroglial clusters plus the general neuronal (excitatory neurons, ExcN; inhibitory neurons, InhN), astrocyte (AS) and microglial (MG) clusters. (F) UMAP plot highlighting in red the most perturbed clusters (6 and 16 in (D)) at 2-month-old, which correspond to MOL and MG, respectively. (G) UMAP plot highlighting in red the most perturbed clusters (O7 and MG in (F)) at 10-month-old, which correspond to MOL and MG, respectively.

**Fig. S7.**
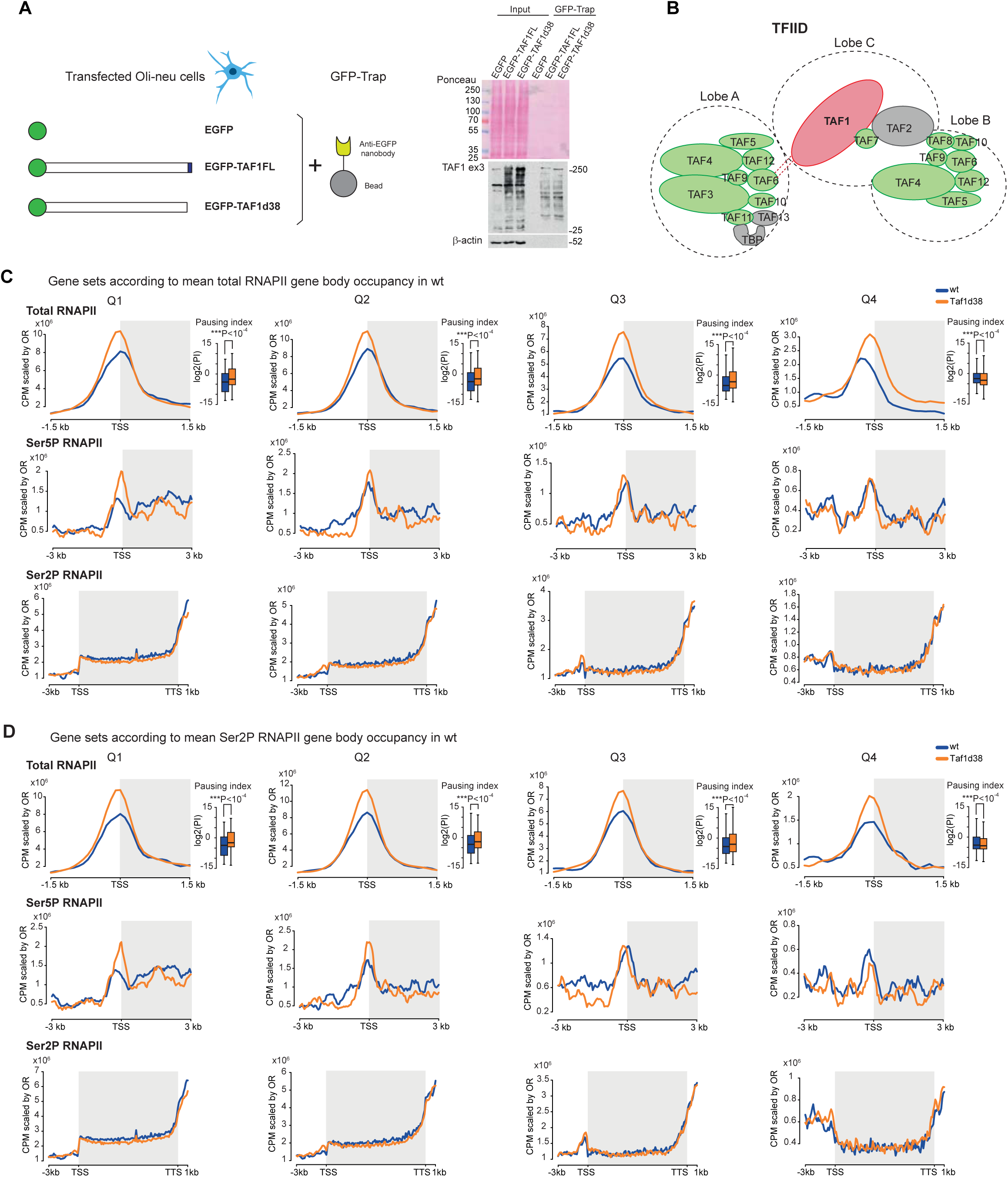
Additional information regarding proteomics in Oli-neu cells and ChIP-seq on mouse samples. (A) Scheme of the GFP-Trap experiment. The anti-EGFP nanobody conjugated to magnetic beads allows capturing EGFP, EGFP-TAF1FL and EGFP-TAF1d38 fusion proteins (left). Enrichment of the bait in TAF1 protein following GFP-Trap immunoprecipitation (right). (B) Scheme of the TFIID subunits that were captured (green) or not captured (gray) after GFP-Trap of EGFP-TAF1FL and EGFP-TAF1d38 samples (modified from Crombie *et al.* 2024). (C) Total RNAPII and Ser5P RNAPII signal at promoters and Ser2P RNAPII signal at gene body (spike-in normalized CPM values, scaled by OR) for gene sets defined according to the quartiles of mean total RNAPII gene body occupancy in wt. The box plots show the distribution of log2(pausing index) in each quartile based on two biological replicates (top). (D) Total RNAPII and Ser5P RNAPII signal at promoters and Ser2P RNAPII signal at gene body (spike-in normalized CPM values, scaled by OR) for gene sets defined according to the quartiles of mean Ser2P RNAPII gene body occupancy in wt. The box plots show the distribution of log2(pausing index) in each quartile based on two biological replicates. (C,D) Mann–Whitney *U* test. Box plots show median ± interquartile range.

